# Design principles of a minimal auxin response system

**DOI:** 10.1101/760876

**Authors:** Hirotaka Kato, Sumanth K. Mutte, Hidemasa Suzuki, Isidro Crespo, Shubhajit Das, Tatyana Radoeva, Mattia Fontana, Yoshihiro Yoshitake, Emi Hainiwa, Willy van den Berg, Simon Lindhoud, Johannes Hohlbein, Jan Willem Borst, D. Roeland Boer, Ryuichi Nishihama, Takayuki Kohchi, Dolf Weijers

## Abstract

Auxin controls numerous growth processes in land plants through a gene expression system that modulates ARF transcription factor activity^1–3^. Gene duplications in families encoding auxin response components have generated tremendous complexity in most land plants, and neofunctionalization enabled various unique response outputs during development^2–4^. However, it is unclear what fundamental biochemical principles underlie this complex response system. By studying the minimal system in *Marchantia polymorpha*, we derive an intuitive and simple model where a single auxin-dependent A-ARF activates gene expression. It is antagonized by an auxin-independent B-ARF that represses common target genes. Expression patterns of both ARF proteins define developmental zones where auxin response is permitted, quantitatively tuned, or prevented. This fundamental design likely represents the ancestral system, and formed the basis for inflated, complex systems.

---

The plant hormone auxin controls essentially all aspects of growth and development, and developmental contexts determine its many unique responses^1,2^. TIR1/AFB F-box proteins perceive auxin and promote ubiquitination and degradation of Aux/IAA transcriptional repressors. Aux/IAAs inhibit DNA-binding ARF transcription factors through direct interaction, and auxin thus releases ARFs from inhibition^3^. Although this signaling module seems simple, each component is encoded by a large gene family in most land plants^4^, allowing overwhelming combinatorial interaction complexity (Fig. 1a; 6 TIR1/AFBs, 29 Aux/IAAs and 23 ARFs; >4000 combinations in *Arabidopsis thaliana*). Given different biochemical properties of family members, sets of response components can trigger unique local responses^2,3^, contributing to the paradoxical functional diversity of the chemically simple auxin hormone. Genetic and functional studies in flowering plants suggest functional interactions and trends in diversification among the many ARFs. ARFs are phylogenetically placed into deeply conserved A/B/C classes^4,5^. A-ARFs are considered transcriptional activators, while some B- and C-ARFs repressors^6^. Systems-wide interaction analysis among Arabidopsis Aux/IAAs and ARFs suggests more prominent auxin-regulation of A-ARFs than B/C-ARFs^7,8^ and individual A- and B-ARFs in the moss *Physcomitrella patens* can antagonize through competition for DNA sites^9^. However, there are several counter-examples where A-ARFs directly repress targets^10^, Aux/IAAs interact with B/C-ARFs^7,8^ and A- and B-ARFs bind different DNA sequences^11^. Because each gene within a multi-gene family may have sub- or neofunctionalized during evolution, it is entirely unclear what basic biochemical architecture underlies the auxin response system. Recently, we have reconstructed the evolutionary history of auxin response components, and found that the irreducible complexity in early-diverging land plants encompasses 1 TIR1/AFB receptor, 1 Aux/IAA and 3 ARFs (A/B/C; Ref. 4). The extant liverwort *Marchantia polymorpha* is representative of this minimal system (Fig. 1a), and may thus resemble the ancestral system before acquiring complexity. Here, we used the *M. polymorpha* auxin response system to understand a minimal auxin response system.

**Fig. 1.**
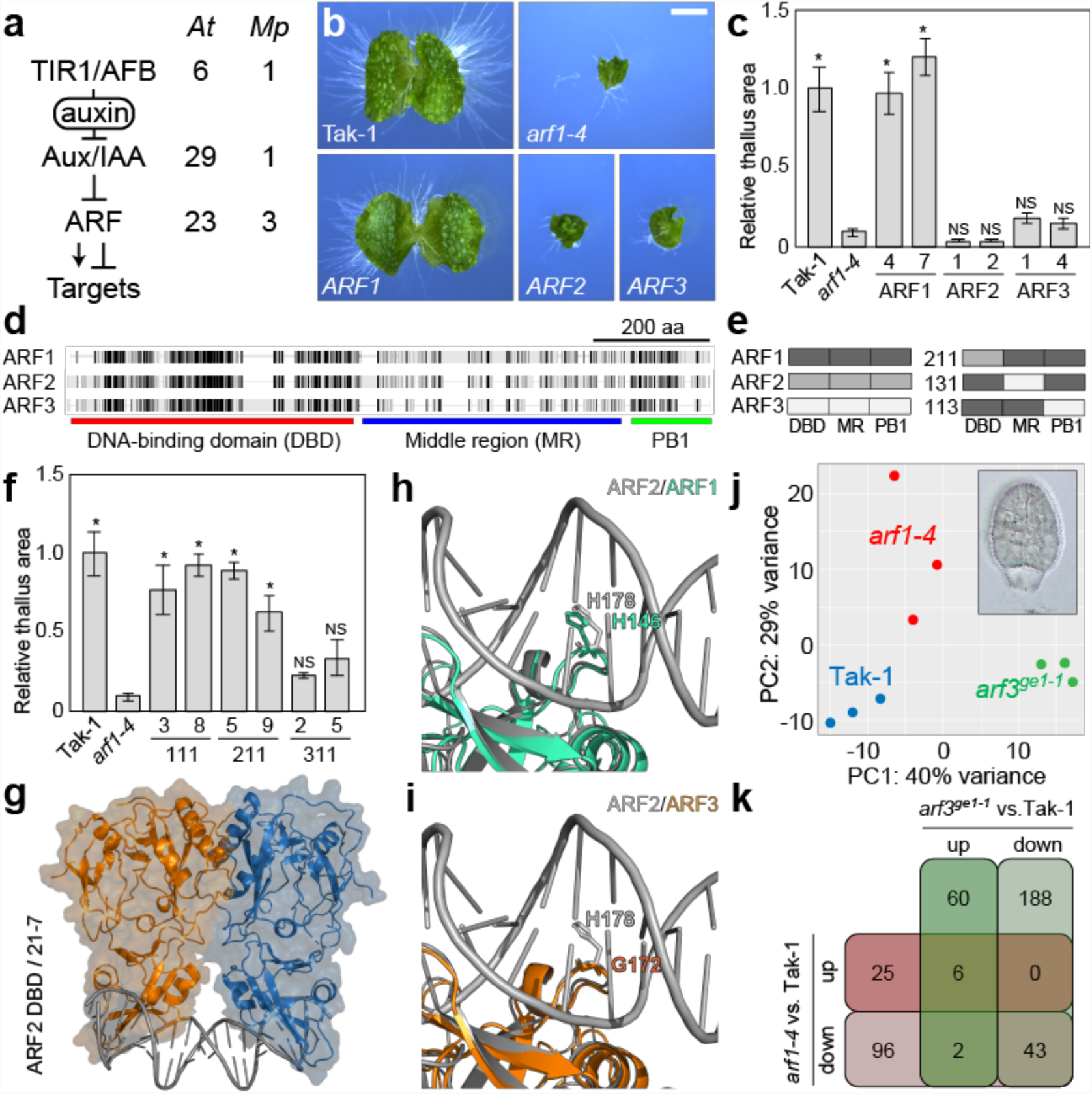
Functional diversity in DNA binding among MpARFs. (**a**) Number of *Arabidopsis thaliana* (*At*) and *Marchantia polymorpha* (*Mp*) auxin response components. (**b**) Ten day-old gemmalings of Tak-1 wild-type, Mp*arf1-4*, _*pro*_Mp*ARF1:*Mp*ARF1* (*ARF1*), _*pro*_Mp*ARF1:*Mp*ARF2* (*ARF2*) and _*pro*_Mp*ARF1:*Mp*ARF3* (*ARF3*), and (**c**) quantification of thallus area in two independent lines for each, normalized to wild-type. (**d**) Similarity and domain organization of MpARFs, and (**e**) nomenclature of domain swaps. (**f**) Relative thallus area in Tak-1 wild-type, Mp*arf1-4*, _*pro*_Mp*ARF1:ARF111* (*111*), _*pro*_Mp*ARF1:ARF211* (*211*) and _*pro*_Mp*ARF1:ARF311* (*311*). **(g)** Crystal structure of MpARF2 DBD in complex with 21-7 DNA at 2.96 Å resolution and structural alignment MpARF2 structure with MpARF1 (**h**) or MpARF3 (**i**) models. Note the lack of conserved His in MpARF3. (**j**) PCA plot of Tak-1, Mp*arf1-4* and Mp*arf3^ge1-1^* gemmae (inset) RNA-seq transcriptomes. (**k**) Overlap of differentially expressed genes in Mp*arf1-4* and Mp*arf3^ge1-1^* gemmae. Bar in (**b**) is 2 mm; error bar in (**c**,**f**) is SD (n=3); asterisks indicate significant differences (p < 0.01) by Tukey-Kramer test; NS: not significant.

Given that functional diversification in *M. polymorpha* is restricted to the ARF family, we first determined if the three *M. polymorpha* ARFs (A: MpARF1, B: MpARF2, C: MpARF3) have unique functions. MpARF1 is a major mediator of auxin-dependent transcription, as mutation causes strong developmental phenotypes, auxin-insensitivity and loss of auxin-dependent transcription^12^. An Mp*arf3* mutant also shows developmental defects, but no evident changes in auxin response^4,13^. When expressed from the Mp*ARF1* promoter, neither MpARF2 nor MpARF3 complemented developmental defects in the Mp*arf1* mutant (Fig. 1b,c), which suggests non-equivalent ARF functions.

ARFs have a typical domain topology with a DNA-binding domain (DBD), a middle region (MR), and a Phox/Bem1 (PB1) oligomerization domain (Fig. 1d). To investigate functional diversification among MpARFs, we performed domain-swaps (Fig. 1e) and expressed chimeric proteins from the Mp*ARF1* promoter in the Mp*arf1* mutant background. ARF111 (each domain derived from MpARF1) fully complemented Mp*arf1* defects (Fig. 1f; **Fig. S1**), suggesting that linkers between domains did not affect function. First, we compared DBD’s functions: ARF211 fully complemented the Mp*arf1* mutant, while ARF311 did not (Fig. 1f; **Fig. S1**). Thus, MpARF2 DBD is functionally equivalent to MpARF1 DBD, while MpARF3 is not. We solved the crystal structure at 2.96 Å resolution (Fig. 1g and **Table S1;** PDB ID: 6SDG) of MpARF2 DBD in complex with a DNA oligonucleotide chosen based on Arabidopsis ARF DNA-binding preference^11^, and identified residues directly contacting DNA (**Fig. S2**). We next generated homology models of MpARF1 and MpARF3 DBDs based on the MpARF2 DBD/DNA co-crystal structure and compared DNA-interacting interfaces. MpARF1 and MpARF2 showed identical DNA interaction interfaces, while MpARF3 deviates at a key position with Gly172 replacing a DNA-contacting histidine (Fig. 1h,i; **Fig. S2**). Thus, structural analysis supports the equivalence of MpARF1 and MpARF2 DNA-binding domains, as well as the non-equivalence of MpARF3 DBD.

To compare *in vivo* target specificity between MpARF1 and MpARF3, we performed transcriptome analysis on Mp*arf1* and Mp*arf3* mutants. Since both are important for gemma development^4,12,13^, we micro-dissected developing gemmae (Fig. 1j; **Fig. S3**) and performed RNA-seq. Principal component analysis (PCA) clustered each genotype separately, suggesting distinct gene expression profiles (Fig. 1j). Furthermore, there was limited overlap between differentially expressed genes in each mutant (Fig. 1k) except for downregulated genes that may reflect impaired gemma development in both mutants. This suggests that MpARF1 and MpARF3 regulate largely different gene sets.

The MR defines the transcriptional activity of ARF proteins, acting as a modular activation and/or repression domain^14^, and MpARF1 and MpARF2 have opposing activities on a model gene when expressed in tobacco^15^. To investigate MR diversity, we replaced the MpARF1 MR with that of MpARF2 or MpARF3 (ARF121, ARF131; a silent mutation was introduced in the MpARF3 MR to prevent microRNA160 regulation^13^). Neither ARF121 nor ARF131 complemented the Mp*arf1* mutant, but instead induced more severe phenotypes, producing callus-like cell masses in both lines (Fig. 2a). These resemble the effects of non-degradable Aux/IAA repressor (mIAA-GR; Fig. 2a; Ref. 15), and suggest active repression. Indeed, while auxin-activated genes were downregulated in Mp*arf1* mutant, they were further repressed in ARF121 and ARF131 plants (Fig. 2b). Despite limited similarity among MRs (Fig. 1d), there is a conserved LFG motif flanked by A/B/C-class-specific residues (Fig. 2d; Ref. 13). The B-ARF-specific (R/K)LFG motif was shown to recruit the TOPLESS (TPL) co-repressor in various transcription factors including Arabidopsis AtARF2 (Ref. 16). Likewise, MpARF2, but not MpARF1, showed clear interaction with MpTPL, which was lost when the LFG motif was substituted by alanines (Fig. 2d; **Fig. S4**). A 17 amino acid fragment encompassing the LFG motif of all MpARFs, but not LFG-to-AAA mutant versions, showed repressor activity when co-expressed with MpTPL in tobacco cells (Fig. 2e; **Fig. S5**). Finally, deletion of the LFG motif in MpARF2 and MpARF3 (ARF12_Δ_1 and ARF13_Δ_1) eliminated the strong phenotypes seen in ARF121 and ARF131 lines, and partially complemented the Mp*arf1* mutant (**Fig. S6a**), yet did not mediate auxin-dependent gene activation (**Fig. S6b**). We could not detect a function for the conserved LFG motif in MpARF1, as deletion did not affect the ability to complement the Mp*arf1* mutant (**Fig. S6a**), or to mediate auxin-dependent gene activation or repression (**Fig. S6c**). Thus, while MpARF1 activates transcription, MpARF2 and MpARF3 recruit MpTPL to repress transcription.

**Fig. 2.**
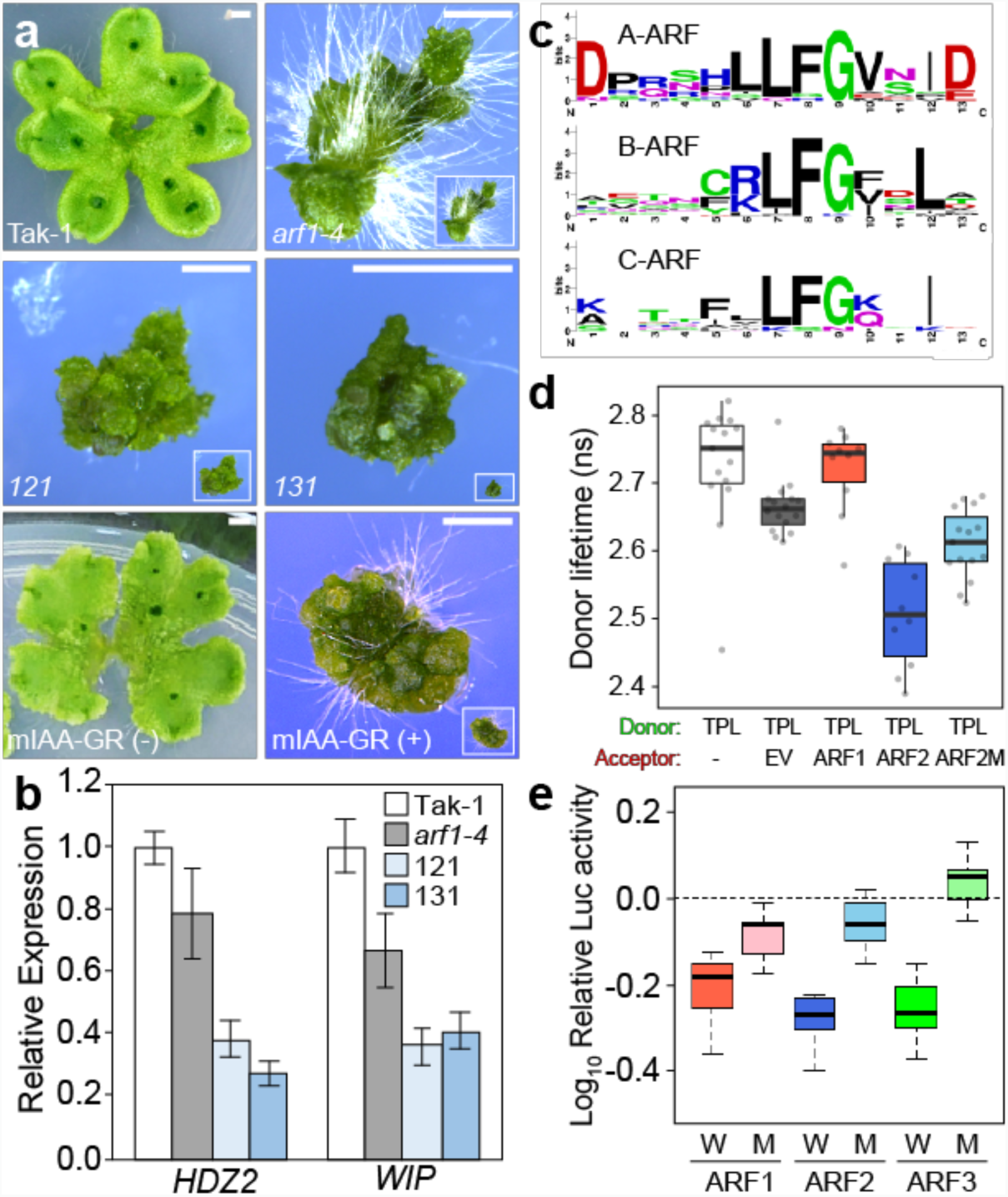
Activation and direct repression by MpARFs. (**a**) Twenty-two day-old gemmalings of Tak-1 wild-type, Mp*arf1-4*, _*pro*_Mp*ARF1:ARF121* (*121*) and _*pro*_Mp*ARF1:ARF131* (*131*) lines, and 20 day-old gemmalings of mIAA-GR grown without (-) or with 1 µM DEX (+). Insets show same magnification as Tak-1. (**b**) Relative qPCR expression of *HDZ2* and *WIP* genes in Tak-1 wild-type, Mp*arf1-4*, ARF*121* and ARF*131*. (**c**) Sequence logos of conserved MR motifs among Marchantia, Physcomitrella, *Selaginella moellendorffii*, *Amborella trichopoda*, and Arabidopsis ARFs. (**d**) FRET-FLIM interaction in Arabidopsis protoplasts: MpTPL-mNeongreen fluorescence lifetime (ns) alone (-) or upon co-expression with mScarlet-I (EV), or fusions to MpARF1, MpARF2 or MpARF2 with LFG→AAA mutation. (**e**) Transcriptional repression of Fluc transgene by co-expression of MpTPL and LFG motif peptides of MpARF1, MpARF2 or MpARF3 (W), or their LFG→AAA mutants (M), relative to a control using Gal4 DBD only in tobacco leaves. Bar in (**a**) = 2 mm.

ARF PB1 domains confer auxin-dependence through interaction with Aux/IAAs^3^, and can also homo-oligomerize^17,18^, *in vivo* significance for which is unknown. PB1 domains were swapped among MpARFs: while ARF112 partially complemented the morphological defects of the Mp*arf1* mutant, ARF113 did not (Fig. 3a,b). Despite complementation under standard conditions, ARF112 did not show any response to auxin (Fig. 3a,b). Thus, MpARF2 and MpARF3 appear to function independently of auxin, which is consistent with limited Arabidopsis B/C-ARF -Aux/IAA interactions^7,8^. To determine the structural basis for potential differential interactions, we generated structural homology models of MpIAA and MpARF1-3 PB1 domains. These showed marked differences in surface charge, mainly on the negative face, (Fig. 3c), which suggests unique interaction properties of each PB1 domain.

**Fig. 3.**
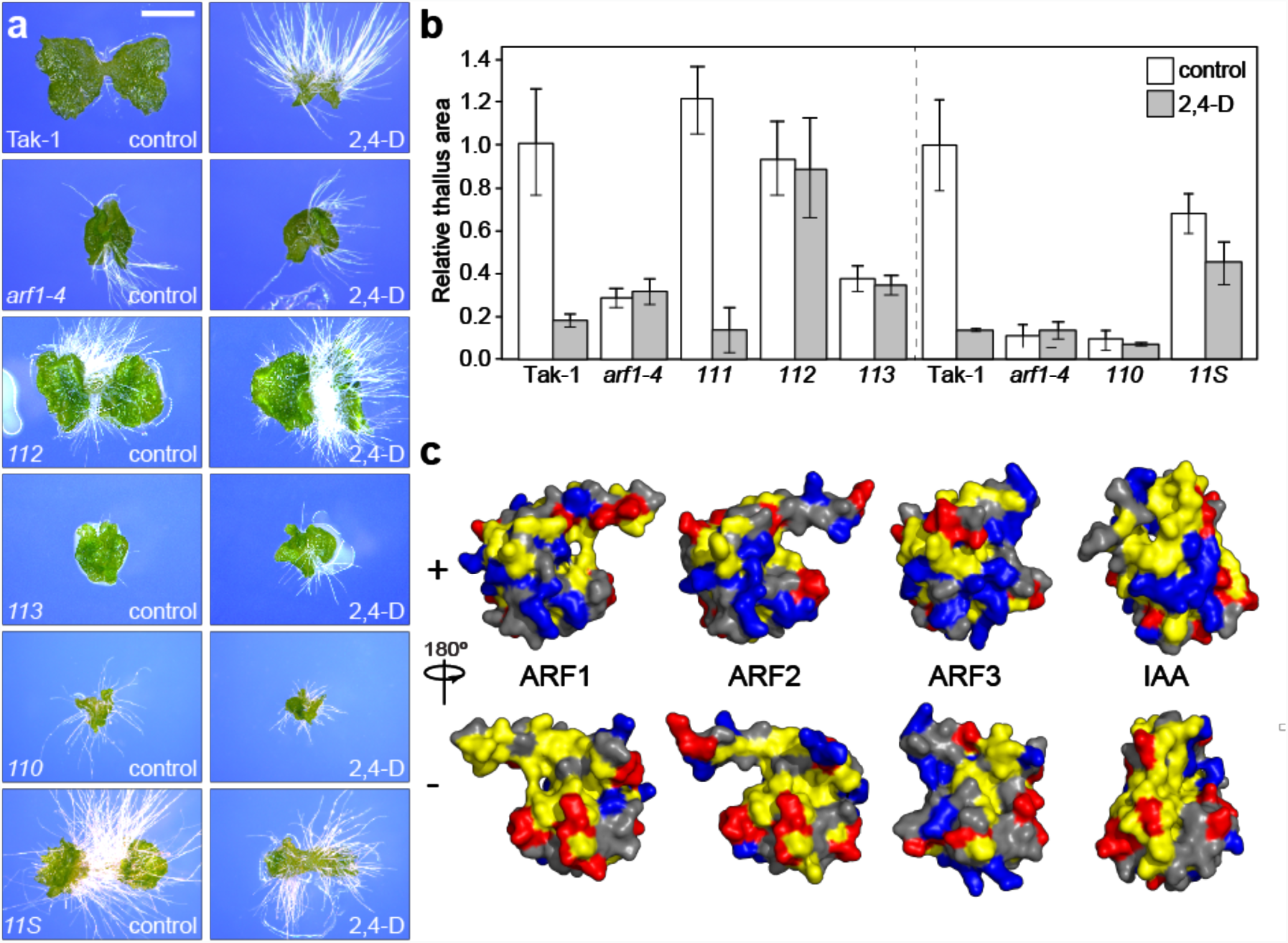
Dual function of the PB1 domain. (**a**) Ten day-old gemmalings of Tak-1 wild-type, Mp*arf1-4*, _*pro*_Mp*ARF1:ARF112* (*112*), _*pro*_Mp*ARF1:ARF113* (*113*), _*pro*_Mp*ARF1:ARF110* (*110*), and _*pro*_Mp*ARF1:ARF11S* (*11S*) lines grown on control medium or on 3 µM 2,4-D, and (**b**) quantification of thallus area relative to Tak-1. Dashed line separates independent experiments. (**c**) Charge distribution (blue, positive; red, negative; yellow, hydrophobic) on surface representations of MpARF1, MpARF2, MpARF3 and MpIAA PB1 domains. Top row shows positive (K) side, and bottom row negative (OPCA) side. Bar in (**a**) = 2 mm; error bar in (**b**) is SD (n=3).

The MpARF2 PB1 domain can replace the MpARF1 PB1 domain under standard, but not auxin-induced conditions (Fig. 3a,b), which suggests function of the MpARF2 PB1 domain is independent of auxin. MpARF1 lacking its PB1 domain (ARF110) could not complement the Mp*arf1* mutant (Fig. 3a,b), suggesting that homo-oligomerization is required for MpARF1 function. To test sufficiency, we replaced the MpARF1 PB1 domain with the Arabidopsis LEAFY SAM oligomerization domain (ARF11S; Ref. 19). ARF11S indeed partially complemented Mp*arf1* mutant defects, producing flat thalli with proper organs (Fig. 3a,b). This demonstrates the importance of homotypic interaction and suggests that this property is shared between A- and B-ARFs, but not C-ARFs, which is consistent with previous interaction assays^15^. Given that MpARF3 is not auxin-regulated and controls different genes from MpARF1, this transcription factor seems unrelated to auxin response in *M. polymorpha*.

From genetic and biochemical experiments, a simple model emerges: A- and B-ARFs compete for the same DNA sites, where the A-ARF can switch from Aux/IAA-mediated repression to auxin-dependent activation, and the B-ARF dampens auxin-responsiveness by direct repression. To test if MpARF2 indeed antagonizes MpARF1, we misexpressed a DEX-inducible MpARF2-GR protein from the constitutive Mp*EF1α* promoter (_*pro*_Mp*EF:*Mp*ARF2-GR*; Ref. 20). As predicted, DEX treatment conferred clear auxin-resistance and rapidly suppressed auxin-dependent gene activation (Fig. 4a,b). We next increased levels of MpARF1, MpARF2 or both. MpARF1 induced growth retardation yet maintained normal auxin sensitivity, while MpARF2 overexpression caused auxin insensitivity (**Fig. S7**). Joint MpARF1/MpARF2 misexpression recovered normal auxin sensitivity (**Fig. S7**), which shows that MpARF2 indeed antagonizes MpARF1, and that MpARF1/2 stoichiometry, rather than absolute MpARF1 or MpARF2 levels, determine auxin responsiveness.

**Fig. 4.**
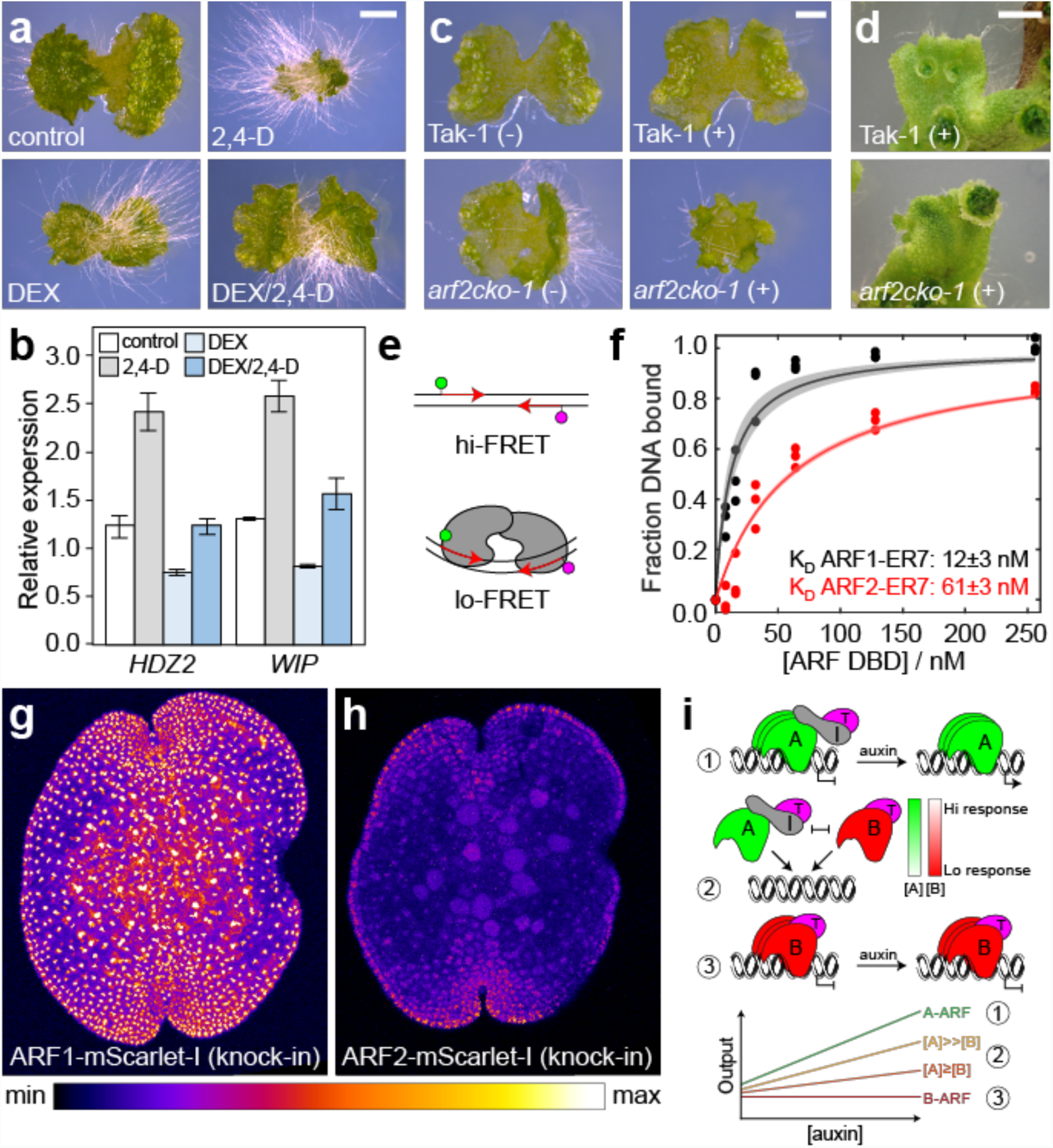
A/B-ARF antagonism defines a minimal auxin response model. (**a**) Ten-day-old gemmalings of a _*pro*_*EF:*Mp*ARF2-GR* line grown on control medium, on 3 µM 2,4-D, on 1 µM DEX, or both. (**b**) Relative qPCR expression of *HDZ2* and *WIP* genes in a _*pro*_*EF:*Mp*ARF2-GR* line treated with control medium, with 10 µM 2,4-D, with 10 µM DEX, or both, for 1 hours. (**c**) Eight-day-old gemmalings and 22-day-old thallus tip (**d**) of Tak-1 or Mp*arf2-1^cko^* grown on control medium without heat shock (-) or with 1 µM DEX and a heat shock (+) on day 1. (**e**) FRET-based assay for measuring ARF binding to inverted repeat ARF binding sites (red arrows). Upon protein binding, FRET efficiency between dyes (magenta, green) changes. (**f**) Quantification of protein-DNA binding, expressed as fraction of bound DNA molecules at varying MpARF1 and MpARF2 DBD concentrations. Lines show fit through raw data within 95% Confidence Interval. (**g**,**h**) Accumulation of MpARF1-mScarlet-I (**g**) and MpARF2-mScarlet-I (**h**) fusion proteins in genomic knock-in lines in mature gemmae in false-color intensity scale. (**i**) Minimal auxin response model: (1) At low [auxin], A-ARF (A) recruits Aux/IAA (I) and TPL (T) to repress genes; at high [auxin], I is degraded, and A activates genes. (2) A and B-ARF (B) compete for binding to the same targets, where stoichiometry of both determines auxin responsiveness. (3) B-ARF recruits TPL to repress targets independently of auxin. Graph: presence and stoichiometry of A and B-ARFs defines cellular responsiveness to auxin. Bars: 1 mm in (**a**,**c**) and 2 mm in (**d**).

To determine the biological significance of the antagonistic MpARF2 activity, we attempted to generate loss-of-function mutants through homologous recombination (HR; Ref. 21) or CRISPR/Cas9 (Ref. 22) with three different sgRNAs. Neither yielded heritable mutations despite screening 770 HR transgenics and 20 chimeric gene-edited lines, suggesting that Mp*ARF2* is an essential gene. We therefore generated Cre/*lox*P-mediated conditional knock-out sectors (Mp*arf2-1^cko^* and Mp*arf2-2^cko^*; Ref. 22,23) through heat-shock and DEX-inducible deletion of a CRISPR-resistant Mp*ARF2* gene in a background with a CRISPR/Cas9-induced mutation in the endogenous Mp*ARF2* gene (**Fig. S8a,b**). Elimination of the additional Mp*ARF2* copy triggered nuclear-localized tdTomato expression, leading to fluorescent mutant sectors (**Fig. S8c,d**). Without induction, Mp*arf2^cko^* mutants produced normal thallus (Fig. 4c; **Fig. S8c**), but Cre-induction caused growth retardation and abnormal morphology in young gemmalings (Fig. 4c; **Fig. S8c**) and growth arrest and gemma cup enlargement in mature thallus (Fig. 4d). These results suggest that MpARF2 is critical for development and meristem maintenance.

Antagonism by direct competition between MpARF1 and MpARF2 is only realistic if their DNA binding affinity for the same DNA target is within the same range, or when there are large differences in protein concentrations. We therefore measured binding affinity of recombinant MpARF1 and MpARF2 DBDs to the same DNA site using a single-molecule FRET-based assay (Fig. 4e; Ref. 24). Proteins were titrated on immobilized DNA oligonucleotide labeled with FRET-compatible fluorophores (**Fig. S9a**), such that protein binding would alter FRET efficiency (**Fig. S9b**), measured at single-molecule resolution (**Fig. S9c**). As DNA, we used the ER7 sequence consisting of two canonical ARF binding sites spaced by 7 nucleotides^11^ (**Fig. S9a**), similar to the one co-crystallized with MpARF2 DBD (Fig. 1g) and that should bind to both proteins based on homology models (Fig. 1h). Using this assay, we derived a K_d_ of 12 nM for MpARF1 DBD, and 61 nM for MpARF2 DBD (Fig. 4f), well within one order of magnitude, and compatible with competition at near-stoichiometric protein concentrations.

Finally, to define true endogenous protein levels and accumulation patterns of MpARF1 and MpARF2, we generated genomic knock-in C-terminal translational fusions to mScarlet-I fluorescent protein. Given that loss- and gain-of function of both Mp*ARF1* and Mp*ARF2* causes strong developmental defects (Fig. 1b; Fig 4c,d; **Fig. S7,8**; Ref. 12), the normal phenotype of these knock-in lines (Fig. 4g,h) demonstrates that these report endogenous protein levels. While MpARF1-mScarlet-I broadly accumulates in gemmae (Fig. 4g), MpARF2-mScarlet-I is restricted to regions surrounding the apical notch (Fig. 4h). Importantly, the spatial distribution patterns of both proteins define distinct zones of cells that express only MpARF1 and MpARF2 in different stoichiometries, and should thus confer differential auxin sensitivity.

Our study reveals a simple model revolving around two competing transcription factors underlying auxin response in the minimal *M. polymorpha* system (Fig. 4i): A-ARF is the only auxin-sensitive transcriptional regulator and can switch from Aux/IAA-mediated repression to activation in an auxin-dependent manner. B-ARF functions independently of auxin and antagonizes A-ARF by competing for target sites and by recruiting the TPL co-repressor. Thus, expression patterns of A-ARF and B-ARF create zones with different auxin sensitivity. It is intuitive how auxin response may have evolved by influencing the stoichiometry of a pre-existing antagonistic ARF pair^25^. This basic module may form the basis for auxin response diversity in species with expanded gene families. Duplication of A- or B-ARFs would change A/B stoichiometry, and also allow duplicated genes to “escape” ancestral regulation, for example by gain of Aux/IAA interaction in B-ARFs^7,8^. Strikingly, the simple model is consistent with most, if not all, studies in the more complex Physcomitrella and Arabidopsis, rationalizing genetic interactions^9^, protein interactions^7,8^, activity assays^6,10^ and *ARF* gene expression patterns^26^, and may therefore represent a universal unit at the base of the complex auxin response networks.

## Acknowledgments

The authors thank the XALOC staff at the synchrotron ALBA and Vanessa Polet Carrillo Carrasco, Shiho Kiryu, Minami Katayama, and Lisa Olijslager for experimental support, Dorus Gadella Kazuyuki Hiratsuka for providing materials, Kimitsune Ishizaki for supporting manuscript preparation, and Ottoline Leyser for comments in the manuscript. This work was supported by an EMBO Long-term Fellowship (ALTF 415-2016) to H.K., a PhD fellowship from the Graduate School Experimental Plant Sciences to J.H. and D.W., a VICI grant (865.14.001) from the Netherlands Organization for Scientific research (NWO) to D.W., the Ministry of Economy and Competitiveness of the Spanish Government [BIO2016-77883-C2-2-P and FIS2015-72574-EXP] (AEI/FEDER,EU) to D.R.B., and ALW-open grant (ALWOP.402) from the Netherlands Organization for Scientific research (NWO) to J.W.B. and JSPS/MEXT KAKENHI (18J12698 to H.S, 19K016166 to Y.Y., 18H04836 to R.N. and 25113009, 15K21758 and 19H05675 to T.K.) and SPIRITS 2017 of Kyoto University to R.N..

## Author contributions

Conceptualization: H.K., R.N., T.K., D.W.; Investigation: H.K., S.K.M., I.C., T.R., S.D., M.F., W.vdB., S.L.; Formal analysis: H.K., S.K.M., I.C., M.F.; Supervision: J.H., R.B., R.N., T.K., D.W.; Funding acquisition: J.H., R.N., T.K., J.W.B., D.W.; Writing – Original Draft: H.K., D.W.; Writing – review & editing: all authors.

## Data and materials availability

All materials generated in this study are freely available upon request to the corresponding author. All data is available in the main text or the supplementary materials. Crystallographic data is available from the RCSB PBD (accession number 6SDG), RNAseq data is available from NCBI Sequence Read Archive (SRA) under the project accession number PRJNA554398 (http://www.ncbi.nlm.nih.gov/bioproject/554398).

## Supplementary Materials

Materials and Methods

References

Figures S1-S9

Tables S1-S2

Supplemental File S1

## References

1. S. Vanneste, J. Friml, Auxin: a trigger for change in plant development. Cell 136, 1005 (2009).

2. Y. Du, B. Scheres, Lateral root formation and the multiple roles of auxin. J. Exp. Bot. 69, 155 (2018).

3. D. Weijers, D. Wagner, Transcriptional responses to the auxin hormone. Annu. Rev. Plant Biol. 67, 539 (2016).

4. S. K. Mutte et al., Origin and evolution of the nuclear auxin response system. eLife 7, e33399 (2018).

5. C. Finet, A. Berne-Dedieu, C. P. Scutt, F. Marlétaz, Evolution of the ARF gene family in land plants: old domains, new tricks. Mol. Biol. Evol. 30, 45 (2013).

6. T. Ulmasov, G. Hagen, T. J. Guilfoyle, Activation and repression of transcription by auxin response factors. Proc. Nat. Acad. Sci. U. S. A. 96, 5844 (1999).

7. S. Piya, S. K. Shrestha, B. Binder, C. N. Stewart, Jr., T. Hewezi, Protein-protein interaction and gene co-expression maps of ARFs and Aux/IAAs in Arabidopsis. Front. Plant Sci. 5, 744 (2014).

8. T. Vernoux et al., The auxin signalling network translates dynamic input into robust patterning at the shoot apex. Mol. Syst. Biol. 7, 508 (2011).

9. M. Lavy et al., Constitutive auxin response in *Physcomitrella* reveals complex interactions between Aux/IAA and ARF proteins. eLife 5, (2016).

10. Z. Zhao et al., Hormonal control of the shoot stem-cell niche. Nature 465, 1089 (2010).

11. D. R. Boer et al., Structural basis for DNA binding specificity by the auxin-dependent ARF transcription factors. Cell 156, 577 (2014).

12. H. Kato et al., The roles of the sole activator-type auxin response factor in pattern formation of *Marchantia polymorpha*. Plant Cell Physiol. 58, 1642 (2017).

13. E. Flores-Sandoval et al., Class C ARFs evolved before the origin of land plants and antagonize differentiation and developmental transitions in *Marchantia polymorpha*. New Phytol. 218, 1612 (2018).

14. S. B. Tiwari, G. Hagen, T. Guilfoyle, The roles of auxin response factor domains in auxin-responsive transcription. Plant Cell 15, 533 (2003).

15. H. Kato et al., Auxin-mediated transcriptional system with a minimal set of components is critical for morphogenesis through the life cycle in *Marchantia polymorpha*. Plos Genet. 11, e1005084 (2015).

16. H. S. Choi, M. Seo, H. T. Cho, Two TPL-binding motifs of ARF2 are involved in repression of auxin responses. Front. Plant Sci. 9, 372 (2018).

17. M. H. Nanao et al., Structural basis for oligomerization of auxin transcriptional regulators. Nat. Commun. 5, 3617 (2014).

18. D. A. Korasick et al., Molecular basis for AUXIN RESPONSE FACTOR protein interaction and the control of auxin response repression. Proc. Nat. Acad. Sci. U. S. A. 111, 5427 (2014).

19. C. Sayou et al., A SAM oligomerization domain shapes the genomic binding landscape of the LEAFY transcription factor. Nat. Commun. 7, 11222 (2016).

20. K. Ishizaki et al., Development of gateway binary vector series with four different selection markers for the liverwort *Marchantia polymorpha*. PLoS One 10, e0138876 (2015).

21. K. Ishizaki, Y. Johzuka-Hisatomi, S. Ishida, S. Iida, T. Kohchi, Homologous recombination-mediated gene targeting in the liverwort *Marchantia polymorpha* L. Sci. Rep. 3, 1532 (2013).

22. S. S. Sugano et al., Efficient CRISPR/Cas9-based genome editing and its application to conditional genetic analysis in *Marchantia polymorpha*. PLoS One 13, e0205117 (2018).

23. R. Nishihama, S. Ishida, H. Urawa, Y. Kamei, T. Kohchi, Conditional gene expression/deletion systems for *Marchantia polymorpha* using its own heat-shock promoter and Cre/*lox*P-mediated site-specific recombination. Plant Cell Physiol. 57, 271 (2016).

24. J. Hohlbein, T. D. Craggs, T. Cordes, Alternating-laser excitation: single-molecule FRET and beyond. Chem. Soc. Rev. 43, 1156 (2014).

25. O. Leyser, Auxin signaling. Plant Physiol. 176, 465 (2018).

26. E. H. Rademacher et al., A cellular expression map of the Arabidopsis AUXIN RESPONSE FACTOR gene family. Plant J. 68, 597 (2011).

