## Supplemental Materials for "Design principles of a minimal auxin response system"

**This PDF file includes:**

Materials and Methods
References
Figs. S1 to S9
Tables S1 to S2

### Materials and Methods

Plant materials and growth conditions

Male *Marchantia polymorpha* strain Takaragaike-1 (Tak-1) was used as wild type. Female strain Tak-2 was used to generate spores for transformation. Cultivation and transformation methods of *M. polymorpha* were described previously[27-29]. *Nicotiana tabacum* were used in transcriptional repression assays. *Arabidopsis thaliana* strain Col-0 was used to generate protoplast for FRET-FLIM experiment.

Promoter and domain swap experiments

Promoter region of MpARF1 including 3 kb upstream from transcriptional start site was amplified with the primer set of HK120/ HK125 (See **Table S2** for all oligonucleotide sequences) and cloned into pMpGWB301 (Addgene) using *Xba*I site (pHKDW046). For promoter swap experiments, genomic coding sequence of MpARFs were amplified with following primer sets; MpARF1: HK009/HK015, MpARF2: HK108/HK119, MpARF3: HK033/HK097, and sub-cloned using pENTR/D-TOPO vector (Thermo Fisher Scientific). Genomic CDS cassettes were then transferred into pHKDW046 using Gateway LR Clonase II Enzyme mix (Thermo Fisher Scientific). DBD of MpARFs were amplified and attached with *Pac*I and *Pme*I restriction enzyme sites by PCR using the primer sets; MpARF1: HK009/HK011, MpARF2: HK018/HK020, MpARF3: HK112/HK113, and then sub-cloned into pENTR/D-TOPO vector. MR of MpARFs with *Pac*I site at 5' end and *Pme*I site at 3' end were amplified from cDNA using the primer sets; MpARF1: HK116/HK120, MpARF1 with stop: HK116/HK285, MpARF2: HK021/HK022, MpARF3: HK030/HK031. PB1 domain of MpARFs or SAM domain of AtLFY with *Pme*I site at 5' end and *Asc*I site at 3' end were amplified from cDNA of *M. polymorpha* or *A. thaliana* using the following primer sets; MpARF1: HK014/HK015, MpARF2: HK023/HK024, MpARF3: HK032/HK033, SAM: HK481/HK482. Amplified MR, PB1, and SAM were then fused with DBD in various combination using the restriction enzyme sites *Pac*I, *Pme*I and *Asc*I. Resultant chimeric CDS were transferred into pHKDW046 with LR reaction to fuse with Mp*ARF1* promoter. All primes used in this study are listed in **Table S2**. At least three independent transformants showing similar phenotype were analyzed for each construct.

Recombinant protein expression and purification

The DNA-binding domains for MpARF1 and MpARF2 used in single-molecule DNA-binding assays and crystallography were amplified using oligonucleotides MpARF1-DBD-FW/ MpARF1-DBD-RV (MpARF1) and MpARF2-DBD-FW/ MpARF2-DBD-RV (MpARF2), and cloned in to a modified pTWIN1 vector[11] by SLiCE cloning[30]. Expression and purification were performed as described for AtARF1 and AtARF5 DNA-binding domains previously[11].

Crystal structural analysis

The oligonucleotide 5'-d(TTGTCGGCGATTCGCCGACAA)-3' was annealed with an equimolar amount of the complementary strand, both purchased from Biomers (Ulm, Germany).

MpARF2:DNA complex was prepared by diluting the required amount of dsDNA in [20 mM Tris (pH8.0), 500 mM NaCl, 20mM DTT] and by subsequently adding this solution to a solution of MpARF2 in the same buffer, to a final 2:1 (protein:dsDNA) stoichiometry and a final MpARF2 concentration of 5 mg/mL. The complex was crystallized by applying the sitting-drop vapordiffusion method using [0.2 M Sodium formate, 20% w/v Polyethylene glycol 3,350] as crystallization buffer at 18°C, in drops of 1 μL + 1 μL (complex:reservoir).

Cube-shaped crystals appeared in less than a week and were frozen within a day of appearance. Cryo-cooling in liquid nitrogen was carried out using a cryo-protecting solution containing reservoir solution supplemented with 10% glycerol. X-ray diffraction experiments were performed at beamline BL13 XALOC (ALBA synchrotron, Spain; Ref. 31). Data were processed using the Global Phasing AutoPROC program[32] applying anisotropic cutoffs[33].

The structure was solved with Phaser[34] using the dimerization domain of AtARF1-DBD (PDB ID: 4LDX). A near complete initial model was obtained with Phenix autobuild[35]. The structure was completed by alternate manual model building with Coot v0.8.9.1 (Ref. 36) and refinement with PHENIX 1.16_3549 (Ref. 35). The refinement statistics and model validation parameters of the crystal structures are given in **Table S1**. Figures were prepared using PyMOL v 2.4.0a0.

Homology modeling

The crystal structure of MpARF2 (PDB ID: 6SDG) was used as template for homology modeling on Swissmodel (https://swissmodel.expasy.org/). MpARF1 and MpARF3 protein sequences and the MpARF2 template were uploaded to Swissmodel server and the modeling was run with default configuration. The global and per-residue model quality were ranked using the QMEAN scoring function (-4.86 for MpARF1 model and -5.92 for MpARF3 model).

Homology models of the PB1 domain were generated using Modeller v9.17 (for MpARF1, MpARF2 and MpIAA; Ref. 37) and Phyre2 web-server (for MpARF3; Ref. 38). Structures of the PB1 domains of AtARF5 (PDB ID: 4CHK) and AtIAA17 (PDB ID: 2MUK) were used as templates for modeling Marchantia ARFs and IAA respectively. In Modeller, 100 random models were generated, where the model with lowest DOPE score was chosen as the best model. In Phyre2, the modeling was performed under 'normal' mode. The obtained homology models were visualized using PyMOL v2.2 (www.pymol.org; Schrödinger LLC, New York, USA). The surface representation of various residues are colored according to their charges and hydrophobicity. Hydrophobic residues (Ala, Gly, Val, Ile, Leu, Phe, Met, and Pro) are colored with yellow, polar residues (Asn, Gln, Thr, Ser, Cys, Tyr, and Trp) are colored with grey, positively charged residues (Arg, Lys, and His) are colored with blue and negatively charged (Glu and Asp) residues are colored with red.

mPS-PI staining for developing gemma

Gemma cups were transversely sectioned into 2-3 pieces by knife. Slices of gemma cups were stained by pseudo-Schiff propidium iodide (PS-PI) staining[39] with some modifications. The tissues incubated in fixative solution (75% ethanol, 25% acetic anhydride) at 4°C for 3 days followed by washing with 80% ethanol. Then, tissues were treated with 1% periodic acid for 1 hour and stained with Schiff reagent containing 0.1 mg/ml propidium iodide for 1 h. Confocal laser scanning microscopy was performed using a Leica TCS-SP5 microscope. Excitation wavelength was 561 nm and the detection range was set from 600 to 700 nm. Z-stack images were obtained with 0.5 μm thickness. Optical section images were generated using Volume Viewer in Fiji software[40].

RNA-seq analysis using developing gemmae

To isolate developing gemmae, thallus tips were grown for 1 week and the youngest gemma cups close to the meristematic notches were collected. Gemma isolation was performed following the protocol for isolating Arabidopsis embryo from ovule with some modifications[41]. CellTrics 50 µm filter (Sysmex) was used to remove debris from the crushed gemma cup mixture. 30 to 80 developing gemmae was isolated by micromanipulator for one sample. RNA was extracted using TRIzol Reagent (Thermo Fisher Scientific) with following modifications. Homogenization was performed at 60°C for 30 minutes. After adding isopropanol, samples were mixed with 1.5 µl of GlycoBlue Coprecipitant (Thermo Fisher Scientific) and incubated at -20°C for overnight. Extracted RNA were treated with RNase-free DNase set (QIAGEN), and purified with RNeasy Minelute Cleanup Kit (QIAGEN). Reverse transcription and cDNA amplification were performed following the published protocol[42]. DNA library for sequencing was prepared with SMARTer ThruPLEX DNA-Seq Kit (Takara Bio), and sequenced with NovaSeq6000 (paired-end 150 bp).

Quality assessment for raw RNA-seq reads was performed using FastQC (www.bioinformatics.babraham.ac.uk/projects/fastqc). Illumina adapters at the ends of the paired reads were cleaned up using TrimGalore (v0.5.0; https://github.com/FelixKrueger/TrimGalore) with the parameters "--stringency 5 --paired --length 100 --clip_R1 30 --clip_R2 30". The cleaned FASTQ reads were mapped onto the *Marchantia polymorpha* genome (v3.1 accessed through Phytozome) using HISAT2 v2.1.0 (Ref. 43) with default parameters with "--dta" as an additional setting. Post-processing of SAM/BAM files was performed using SAMTOOLS v1.9 (Ref. 44). FeatureCounts v1.6.2 (Ref. 45) was used to count the raw reads corresponding to each gene, with the parameters "-t 'exon' -g 'gene_name' -Q 30 -p --primary". DEseq2 (Ref. 46) was used to normalize the raw counts and perform the differential expression analysis (Padj < 0.05). All the sequenced raw reads were deposited in NCBI Sequence Read Archive (SRA) under the project accession number PRJNA554398 (http://www.ncbi.nlm.nih.gov/bioproject/554398).

Quantitative RT-PCR

Auxin treatment on 10-day-old gemmalings, RNA extraction, cDNA synthesis, and quantitative RT-PCR were performed as previously described[4]. DEX treatment was similarly performed as auxin treatment.

FRET-FLIM

For FRET-FLIM, fusions of MpTPL were made in pMON999-mNeonGreen, and fusions of MpARF1, MpARF2 and the LFG-AAA mutant of MpARF2 were made in pMON999-mScarlet-I vectors using primers listed in **Table S2** and using SLiCE cloning[30].

Fluorescent proteins were co-expressed in Arabidopsis leaf mesophyll protoplasts, and FLIM analysis was performed as described previously[47]. Measurements were conducted on a Leica TCS SP8 X system equipped with hybrid detectors (HyD) and a pulsed white-light laser. The system was used for detection of mNeonGreen and mScarlet-I, which were excited at 488 nm and 561 nm, and detected between 520-551 nm and 585-607 nm, respectively. From the fluorescence intensity images, the decay curves were calculated per pixel and were fitted with a double-exponential decay model using the SPCImage (Becker & Hickl; version 7.1). For all samples, a double-exponential model function was used without fixing any parameter.

Transcriptional repression assay

The GAL4 DBD sequence was amplified with the primer set (GAL4BD_RwithXbaI/ GAL4BD_FwithXbaI) and cloned into the *Xba*I site of pMpGWB102 (Addgene) to generate pMpGWB102-GAL4DBD that contains a cassette for cauliflower mosaic virus 35S promoter:GAL4DBD-GATEWAY:Nos terminator. The LFG motif sequences were amplified with partially overlapping primer pairs (**Table S2**), and subcloned into the pENTR/D-TOPO. The generated LFG motifs were transferred into pMpGWB102-GAL4DBD using LR Clonase II enzyme mix. MpTPL cDNA was amplified from Tak-1 thallus cDNA with the primer set (MpTPL_F/MpTPL_R1), subcloned into the pENTR/D-TOPO, and transferred into the pMpGWB102 using LR Clonase II enzyme mix. The *Renilla* luciferase coding sequence and the reporter construct (35S enhancer-6xUAS-35S minimal promoter:firefly luciferase) were amplified with the primer sets (RenillaLUCforTOPOF/RenillaLUCstopR and 6xUASfLUCcloningFforTOPO/6xUASfLUC_stopR, respectively), subcloned into the pENTR/D-TOPO, and transferred into pMpGWB102 and pMpGWB101 using LR Clonase II enzyme mix. Plasmids used as templates for amplification of Gal4 DBD, *Renilla* luciferase, and the reporter construct were described previously[48].

*Nicotiana tabacum* plants were grown under continuous white light at 22°C for 4-5 weeks. *Agrobacterium* harboring effector, co-effector, reporter, or internal control vectors were cultured in liquid LB media containing selection antibiotics at 28°C. Mixture of the four cultures (26:15:30:4) were centrifugated at 2,600 g for 5 min, re-suspended in MES buffer (10 mM $MgCl_2$, 10 mM MES, 200 μM acetosyringone) to an OD600 of 1.0, and incubated at 28°C for 4 h before infiltration. Infiltration was performed according to the methods described previously[49]. A set of the *Agrobacterium* mixtures was infiltrated at 8-9 points, and transient expression was assayed 3 days after inoculation. Firefly-luciferase (Fluc) and Renilla-luciferase (Rluc) activities were assayed using the dual-luciferase assay reagents (Promega), and microplate reader POWERSCAN4 (DS PHARMA BIOMEDICAL). 3 days after the inoculation, an approximately 5x5 mm leaf disc was removed and ground in 200 μL of passive lysis buffer. 20 μL of lysate supernatant was incubated with 100 μL of the Luciferase Assay Reagent, and chemiluminescence was measured. After the first measurement, 100 μL of the Stop & Glo Reagent was added, and a second measurement was taken. Relative luciferase activity of each effector condition was calculated as the ratio between the Fluc and the Rluc (standard) activity and divided by that of the control effector condition (Gal4 DBD without LFG motif) from a same leaf to normalize differences among leaves.

Overexpression of MpARF1 and MpARF2

To generate $_{pro}$*EF:MpARF2-GR*, genomic CDS of Mp*ARF2* was amplified with the primer set (HK108/HK109), sub-cloned in pENTR/D-TOPO vector, and then transferred into pMpGWB113 to insert between MpEF1α promoter and GR[20]. To generate $_{pro}$*EF:MpARF1*, MpARF1 cDNA was amplified by RT-PCR using mRNA from Tak-1 as template with the primer set (ARF1CDS_inf_F1/ARF1CDS_inf_R1), cloned into pENTR-1A vector using In-Fusion HD Cloning Kit (TaKaRa), and then transferred to pMpGWB303. To generate $_{pro}$*EF:MpARF2*, MpARF2 cDNA was amplified as for MpARF1 with the primer set (MpARF2_c1f_TOPO/MpARF2_stop), cloned into pENTR/D-TOPO vector, and then transferred to pMpGWB103.

Accessible Volume simulations

The web server 3D-DART[50] was used to model a standard B-DNA structure for the oligo used in the smFRET experiments and to bend it using the geometrical information extracted from the short DNA present in the crystal structure of MpARF2. The standard B-DNA oligo and the bent oligo bound to MpARF2 were then used as starting point to model the accessible volumes of the FRET pair. The accessible volumes were calculated with FRET-restrained positioning and screening (FPS) software[51] using the geometric characteristic for the dyes reported in previously[52].

Single-molecule measurement of dissociation constant

Imaging was performed on a home-built TIRF microscope as previously described[53]. The measurements were performed using alternating-laser excitation (ALEX); in this excitation scheme a frame where the donor is excited is followed by a frame where the acceptor is excited and so on. The emission of the fluorophores is spectrally divided into two different channels on the emCCD camera sensor. This creates three photon streams, donor emission after donor excitation (DD), acceptor emission after donor excitation (DA, arising from FRET) and acceptor emission after acceptor excitation (AA). These photon streams can be used to calculate the raw FRET efficiency ($E^*=DA/(DD+DA)$) and stoichiometry ($S=(DD+DA)/(DD+DA+AA)$). $E^*$ contains the information about the relative distance of the two fluorophores whereas S contains information about the labelling state of a given molecule (allowing to filter out molecules missing an active donor or an active acceptor).

The camera acquisition time and the excitation time were set to 250 ms; laser powers were set to 0.5 mW for both green ($\lambda$ = 561 nm) and red ($\lambda$ = 638 nm) lasers. The imaging buffer contained 137 mM NaCl, 2.7 mM KCl, 10 mM phosphate, 1 mM Trolox, 1% gloxy and 1% glucose. Trolox is a triplet state quencher[54]; gloxy is an enzymatic system that uses glucose as substrate for oxygen removal, decreasing fluorophore bleaching[55].

Single-molecule titrations

Labelled dsDNA oligos were immobilized on a PEGylated glass coverslip as described previously[56]. In particular, the PEGylation was carried out inside the wells of a silicone gaskets placed on the coverslip (Grace Bio-labs). Each titration was performed using a single well, washing it between data points with ~600 µL of PBS 1X. The final washing step consisted of three washings separated by 15 minutes. Each data point consisted of three movies (1000 frames each); to allow the system to equilibrate a waiting step of 5 minutes was added before starting the acquisition of the first movie.

Mutagenesis of MpARF2

For mutagenesis of MpARF2 by homologous recombination, homologous arms were amplified from MpARF2 genomic DNA by using the primer set (MpARF2-gt_F1/ MpARF2-gt_R1, MpARF2-gt_F2/ MpARF2-gt_R2) and cloned into pJHY-TMp1 (Ref. 21). Resultant plasmid was transformed into F1 spores generated by crossing Tak-1 to Tak-2 through the Agrobacterium-mediated method (*28*). To check the recombination, genomic PCR was performed using the primer sets (MpARF2-L2/MpEFp_GT_R1, MpARF2-L6/MpARF2-R12, MpARF2-R1/HIF).

For mutagenesis of MpARF2 by CRISPR/Cas9 method, sets of two oligo DNA (sgRNA1: HK184/HK185, sgRNA2: HK186/HK187, sgRNA3: HK188/HK189) were annealed and cloned in pMpGW_En03 (Ref. 22). The resultant cassettes expressing sgRNA and Cas9 were transferred into pMpGE010 by LR reaction. To generate sgRNA-resistant MpARF2, cDNA sequence including CDS was amplified with the primer set (HK119/HK282) and cloned into pENTR/D-TOPO vector. The target site of sgRNA1 was mutated by PCR with the primer set (HK279/HK280) and self-ligation. MpARF2 promoter region including 3 kb upstream of transcriptional start site was also amplified with the primer set (HK107/HK281) and cloned in pENTR/D-TOPO vector. MpARF2 promoter and sgRNA-resistant CDS was combined using the *EcoR*I and *Asc*I sites. The resultant cassette was transferred into pMpGWB337 by LR reaction[23]. To check the mutation in MpARF2 locus, genomic PCR was performed with the primer set (HK283/HK284) which bind introns and thus amplifies only endogenous MpARF2 copy. To delete sgRNA-resistant MpARF2 cassette from the mutant genome, mutants were planted on the medium with 1 μM DEX, incubated at 37°C for 1 h after 1 day cultivation, and then placed back to the normal growth condition. The primer set (HK064/HK190) was used to check the deletion of ARF2m by genomic PCR.

Genomic knock-in translational fusions for MpARF1 and MpARF2

Two homologous arms of ~3.6 kb length each were amplified by PCR from genomic DNA. The 3' end of the genomic CDS excluding the stop codon was amplified using oligonucleotides SJD005/SJD006 (MpARF1) or SJD009/SJD010 (MpARF2), another arm from the noncoding region after stop codon was amplified using SJD007/SJD008 (MpARF1) or SJD011/SJD012 (MpARF2). These fragments were cloned into the pJHY-TMp1 vector[21] at *Hind*III and *Asc*I sites respectively using the SLiCE cloning method[30]. The fluorescent protein mScarlet-I was amplified using oligonucleotides SJD003/SJD004 and cloned downstream of the first homologous arm to be translationally fused with MpARF1/2. Transformation into spores was performed as previously described[28]. Successfully knocked-in lines were screened from 6 (MpARF1) and 78 (MpARF2) hygromycin-resistant transformants by genomic PCR using the primer sets (MpARF1: HK056/SJD091, HK050/SJD149; MpARF2: HK072/SJD094, HK063/HK149).

Live cell confocal imaging was performed on mature gemmae using a Leica SP8X-SMD confocal microscope equipped with a hybrid (HyD) detector and a pulsed white-light laser. mScarlet-I was excited with 561 nm laser line (10 % laser output of WLL) and fluorescence was detected by the hybrid detector from 570-620 nm. Images were acquired in photon counting mode and accumulation of 10 frames per image was taken. To remove auto-fluorescence, time-gating was active in order to suppress auto-fluorescence. Maximum intensity projections of the images were obtained from z-stack series, capturing z-slices with 1.0 um interval and images were processed using Image J.

**Supplemental references**


27. S. Chiyoda, K. Ishizaki, H. Kataoka, K. T. Yamato, T. Kohchi, Direct transformation of the liverwort Marchantia polymorpha L. by particle bombardment using immature thalli developing from spores. *Plant Cell Rep.* **27**, 1467 (2008).
28. K. Ishizaki, S. Chiyoda, K. T. Yamato, T. Kohchi, Agrobacterium-mediated transformation of the haploid liverwort Marchantia polymorpha L., an emerging model for plant biology. *Plant Cell Physiol.* **49**, 1084 (2008).
29. A. Kubota, K. Ishizaki, M. Hosaka, T. Kohchi, Efficient Agrobacterium-mediated transformation of the liverwort Marchantia polymorpha using regenerating thalli. *Biosci. Biotechnol. Biochem.* **77**, 167 (2013).
30. Y. Zhang, U. Werling, W. Edelmann, SLiCE: a novel bacterial cell extract-based DNA cloning method. *Nucleic Acids Res.* **40**, e55 (2012).
31. J. Juanhuix *et al.*, Developments in optics and performance at BL13-XALOC, the macromolecular crystallography beamline at the ALBA synchrotron. *J. Synchrotron Radiat.* **21**, 679 (2014).
32. C. Vonrhein *et al.*, Data processing and analysis with the autoPROC toolbox. *Acta Crystallogr. D Biol. Crystallogr.* **67**, 293 (2011).
33. I. J. Tickle *et al.*, *STARANISO (http://staraniso.globalphasing.org/cgi-bin/staraniso.cgi)*. Cambridge, United Kingdom: Global Phasing Ltd., (2018).
34. A. J. McCoy *et al.*, Phaser crystallographic software. *J. Appl. Crystallogr.* **40**, 658 (2007).
35. P. D. Adams *et al.*, PHENIX: a comprehensive Python-based system for macromolecular structure solution. *Acta Crystallogr. D Biol. Crystallogr.* **66**, 213 (2010).
36. P. Emsley, B. Lohkamp, W. G. Scott, K. Cowtan, Features and development of Coot. *Acta Crystallogr. D Biol. Crystallogr.* **66**, 486 (2010).
37. B. Webb, A. Sali, Comparative Protein Structure Modeling Using MODELLER. *Curr. Protoc. Bioinformatics* **54**, 5 6 1 (2016).
38. L. A. Kelley, S. Mezulis, C. M. Yates, M. N. Wass, M. J. Sternberg, The Phyre2 web portal for protein modeling, prediction and analysis. *Nat. Protoc.* **10**, 845 (2015).
39. E. Truernit, K. R. Siemering, S. Hodge, V. Grbic, J. Haseloff, A map of KNAT gene expression in the Arabidopsis root. *Plant Mol. Biol.* **60**, 1 (2006).
40. J. Schindelin *et al.*, Fiji: an open-source platform for biological-image analysis. *Nat. Methods* **9**, 676 (2012).
41. M. T. Raissig, V. Gagliardini, J. Jaenisch, U. Grossniklaus, C. Baroux, Efficient and rapid isolation of early-stage embryos from Arabidopsis thaliana seeds. *J. Vis. Exp.* **76**, e50371 (2013).
42. J. J. Trombetta *et al.*, Preparation of Single-Cell RNA-Seq Libraries for Next Generation Sequencing. *Curr. Protoc. Mol. Biol.* **107**, 4 22 1 (2014).

43. D. Kim, B. Langmead, S. L. Salzberg, HISAT: a fast spliced aligner with low memory requirements. *Nat. Methods* **12**, 357 (2015).

44. H. Li *et al.*, The Sequence Alignment/Map format and SAMtools. *Bioinformatics* **25**, 2078 (2009).

45. Y. Liao, G. K. Smyth, W. Shi, featureCounts: an efficient general purpose program for assigning sequence reads to genomic features. *Bioinformatics* **30**, 923 (2014).

46. M. I. Love, W. Huber, S. Anders, Moderated estimation of fold change and dispersion for RNA-seq data with DESeq2. *Genome Biol.* **15**, 550 (2014).

47. A. Freire-Rios, T. Radoeva, B. De Rybel, D. Weijers, J.W. Borst. FRET-FLIM for visualizing and quantifying protein interactions in live plant cells. *Meth. Mol. Biol.* **1497,** 135 (2017).

48. N. Matsuo, M. Minami, T. Maeda, K. Hiratsuka, Dual Luciferase Assay for Monitoring Transient Gene Expression in Higher Plants. *Plant Biotechnol.* **18**, 71 (2001).

49. T. Akagi, A. Ikegami, K. Yonemori, DkMyb2 wound-induced transcription factor of persimmon (Diospyros kaki Thunb.), contributes to proanthocyanidin regulation. *Planta* **232**, 1045 (2010).

50. M. van Dijk, A. M. Bonvin, 3D-DART: a DNA structure modelling server. *Nucleic Acids Res.* **37**, W235 (2009).

51. S. Kalinin *et al.*, A toolkit and benchmark study for FRET-restrained high-precision structural modeling. *Nat. Methods* **9**, 1218 (2012).

52. T. D. Craggs *et al.*, Substrate conformational dynamics drive structure-specific recognition of gapped DNA by DNA polymerase. *bioRxiv*, 263038 (2018).

53. S. Farooq, J. Hohlbein, Camera-based single-molecule FRET detection with improved time resolution. *Phys. Chem. Chem. Phys.* **17**, 27862 (2015).

54. T. Cordes, J. Vogelsang, P. Tinnefeld, On the mechanism of Trolox as antiblinking and antibleaching reagent. *J. Am. Chem. Soc.* **131**, 5018 (2009).

55. I. Rasnik, S. A. McKinney, T. Ha, Nonblinking and long-lasting single-molecule fluorescence imaging. *Nat. Methods* **3**, 891 (2006).

56. G. W. Evans, J. Hohlbein, T. Craggs, L. Aigrain, A. N. Kapanidis, Real-time single-molecule studies of the motions of DNA polymerase fingers illuminate DNA synthesis mechanisms. *Nucleic Acids Res.* **43**, 5998 (2015).

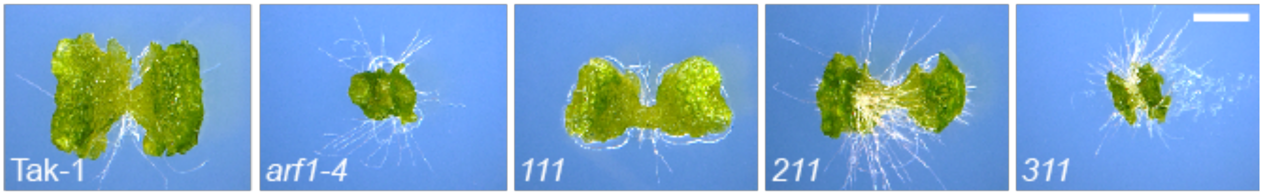


**Fig. S1: Phenotypes of DBD swap lines.**

Ten-day-old gemmalings of Tak-1 wild-type, Mp*arf1-4*, $_{pro}$Mp*ARF1:ARF111* (*111*), $_{pro}$Mp*ARF1:ARF211* (*211*) and $_{pro}$Mp*ARF1:ARF311* (*311*). Bar = 2 mm.

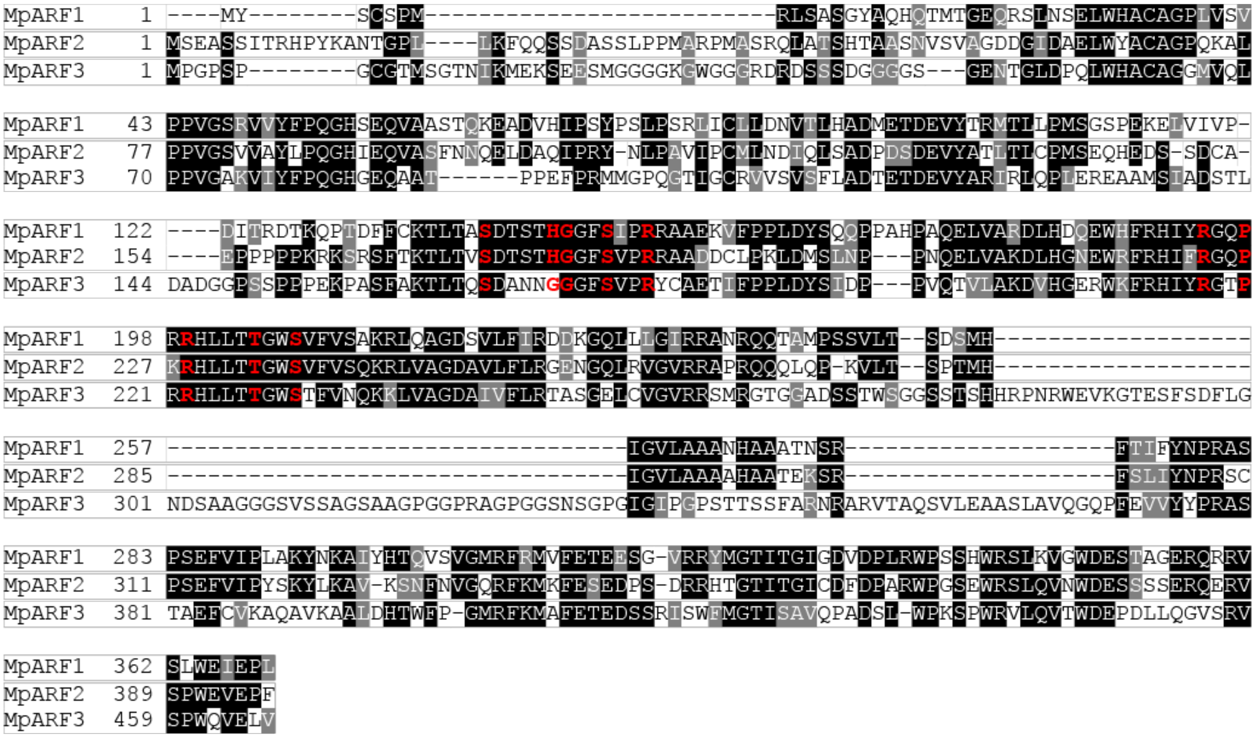


**Fig. S2: Alignment of *M. polymorpha* ARF DBDs.**

Sequence alignment of DNA-binding domains in MpARF1, MpARF2 and MpARF3, delimited by the fragment used for determining the MpARF2 DBD crystal structure. DNA-contacting residues in the MpARF2 crystal structure and the homologous residues in MpARF1 and MpARF3 are indicated in red. Conserved residues are boxed in black, and conservative substitutions are boxed in grey.

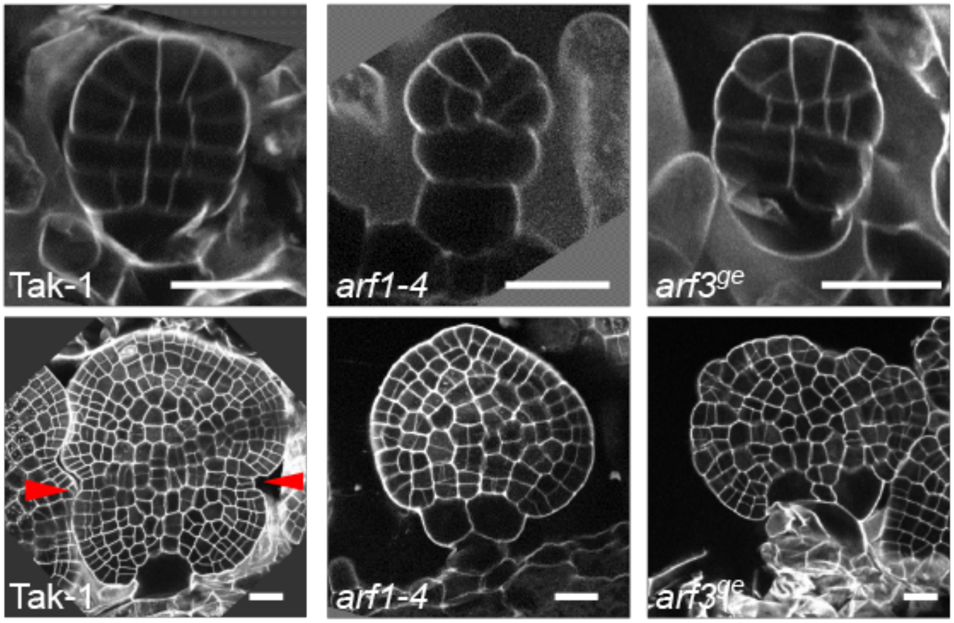


**Fig. S3. Gemmae phenotypes in Mp*arf1* and Mp*arf3* mutants.**

Optical medial sections through early- (top row) and late-stage (bottom row) gemmae from Tak-1, Mp*arf1-4* and Mp*arf3$^{ge1}$-1* plants. Medial sections were taken from full 3D stacks of gemmae in which cell walls were labeled using mPS-PI staining. The positions of the future apical notches are indicated by red arrowheads in Tak-1. Note that these indentations are missing in both mutants. Bars are 20 µm in all panels.

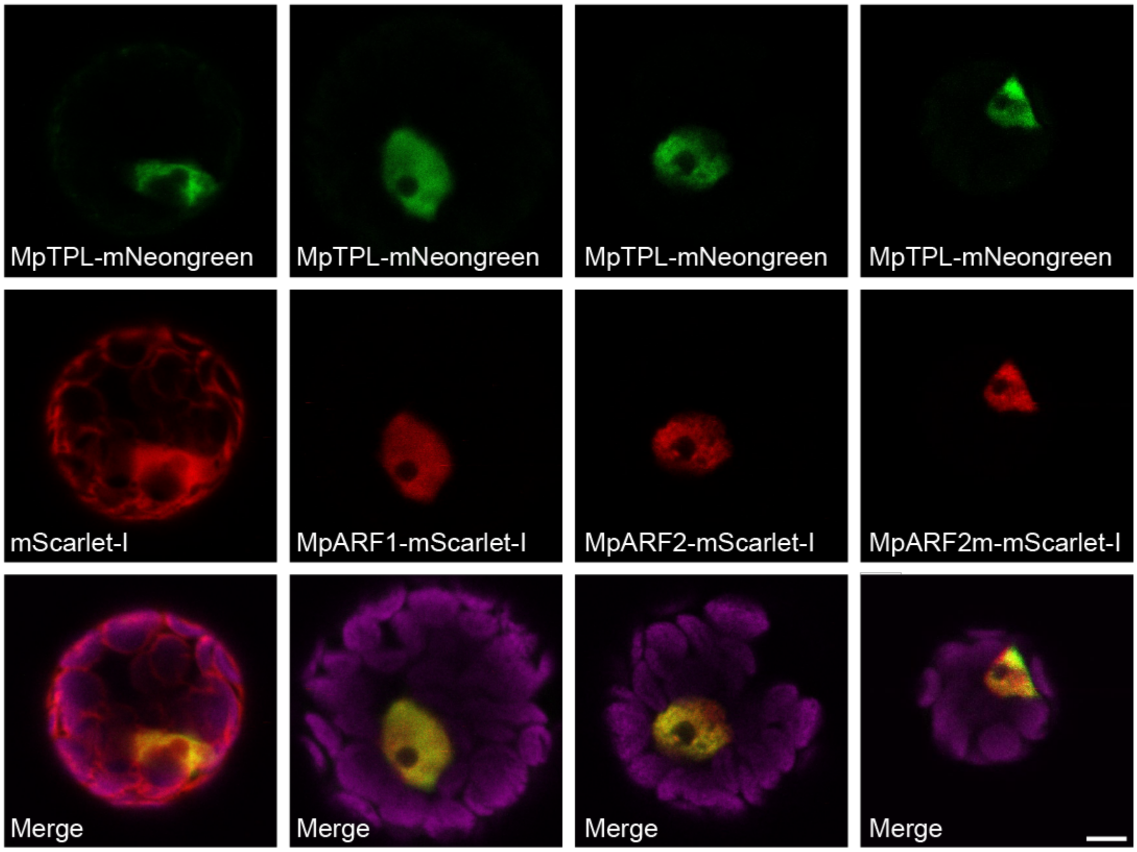


**Fig. S4. Expression of MpTPL and MpARFs for FRET-FLIM**

Fluorescence of MpTPL-mNeongreen (FRET donor; top row), either co-expressed with fusions of mScarlet-I (FRET acceptor; middle row), to empty vector or with MpARF1-mScarlet-I, MpARF2-mScarlet-I or MpARF2(LFG-AAA)-mScarlet-I. Bottom panel shows overlay of mNeongreen, mScarlet-I and chloroplast autofluorescence signals. Note that all fusion proteins localize to nuclei. Bar is 5 µm.

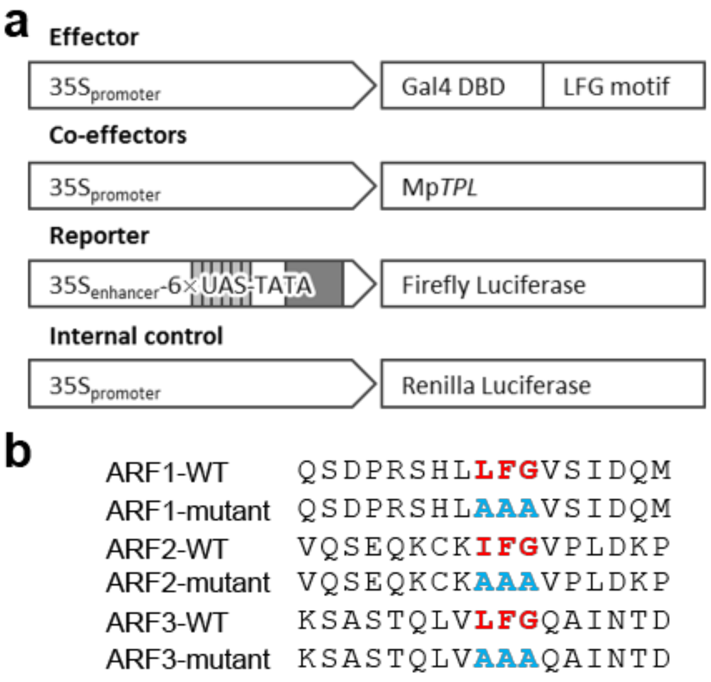


**Fig. S5. Transcriptional repression assay.**

(**a**) Plasmids used in the transcriptional activity assay in *Nicotiana tabacum* leaf. The effector plasmid expresses a fusion between the GAL4 DNA-binding domain (DBD) and the extended LFG motif from wild-type or mutant MpARFs (see **b**) from the *35S* promoter. Mp*TPL* is co-expressed from the *35S* promoter. The reporter plasmid has *Firefly Luciferase* expressed from a minimal *35S* promoter with 6 GAL4-dependent *Upstream Activating Sequence* (*UAS*) elements and a *35S enhancer* sequence. *Renilla Luciferase* is expressed from the *35S* promoter as an internal normalization control. (**b**) Sequences of the wild-type (WT) and mutant peptides from MpARF1, MpARF2 and MpARF3, encompassing the LFG motif.

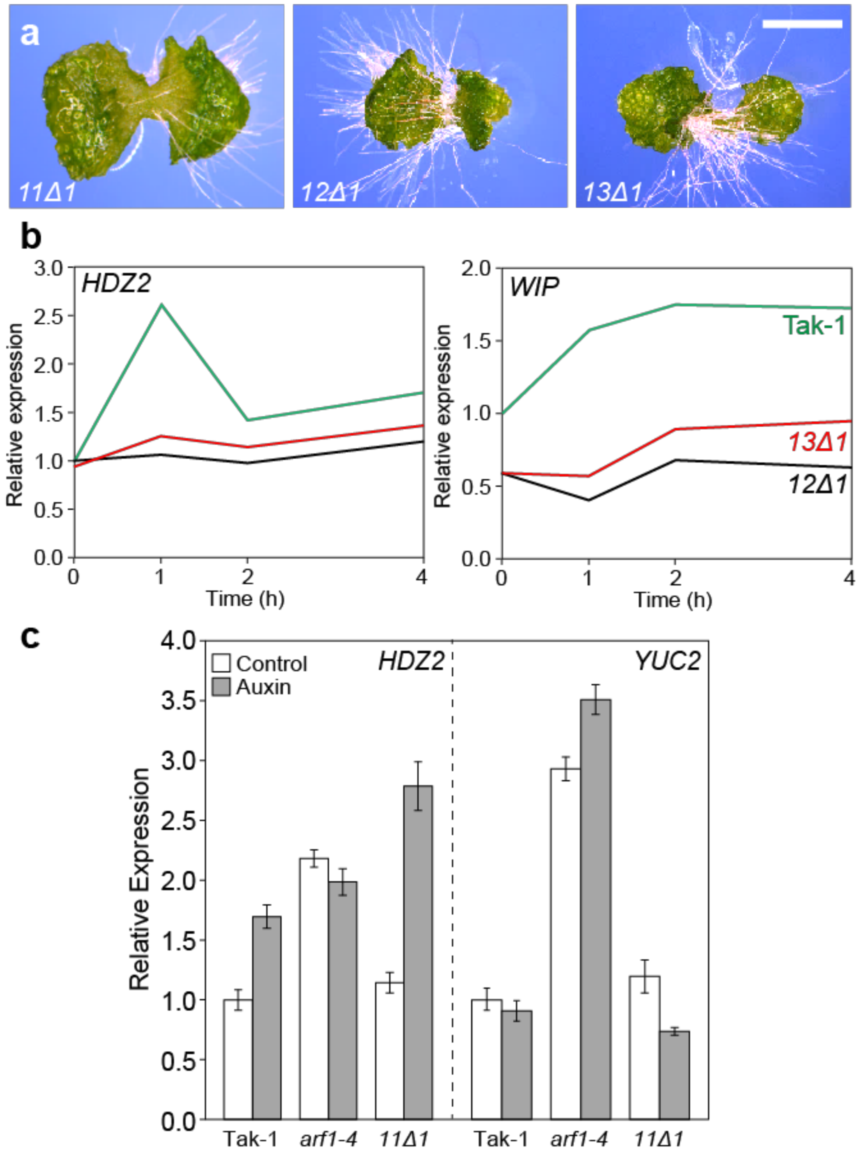


**Fig. S6. Biological significance of the MpARF LFG motifs.**

(**a**) Ten-day-old gemmalings of $_{pro}$Mp*ARF1:ARF11Δ1* (*11Δ1*), $_{pro}$Mp*ARF1:ARF12Δ1* (*12Δ1*) and $_{pro}$Mp*ARF1:ARF13Δ1* (*13Δ1*) lines. (**b**) Relative qPCR expression of *HDZ2* and *WIP* genes in Tak-1 wild-type, $_{pro}$Mp*ARF1:ARF12Δ1* (*12Δ1*) and $_{pro}$Mp*ARF1:ARF13Δ1* (*13Δ1*) gemmalings, 0, 1, 2 and 4 hours after treatment with 10 µM 2,4-D. (**c**) Relative qPCR expression of *HDZ2* and *YUC2* genes in Tak-1 wild-type, Mp*arf1-4* and $_{pro}$Mp*ARF1:ARF11Δ1* (*11Δ1*) gemmalings, after a 1-hour treatment with control media or with 10 µM 2,4-D. Bar in (**a**) = 2 mm.

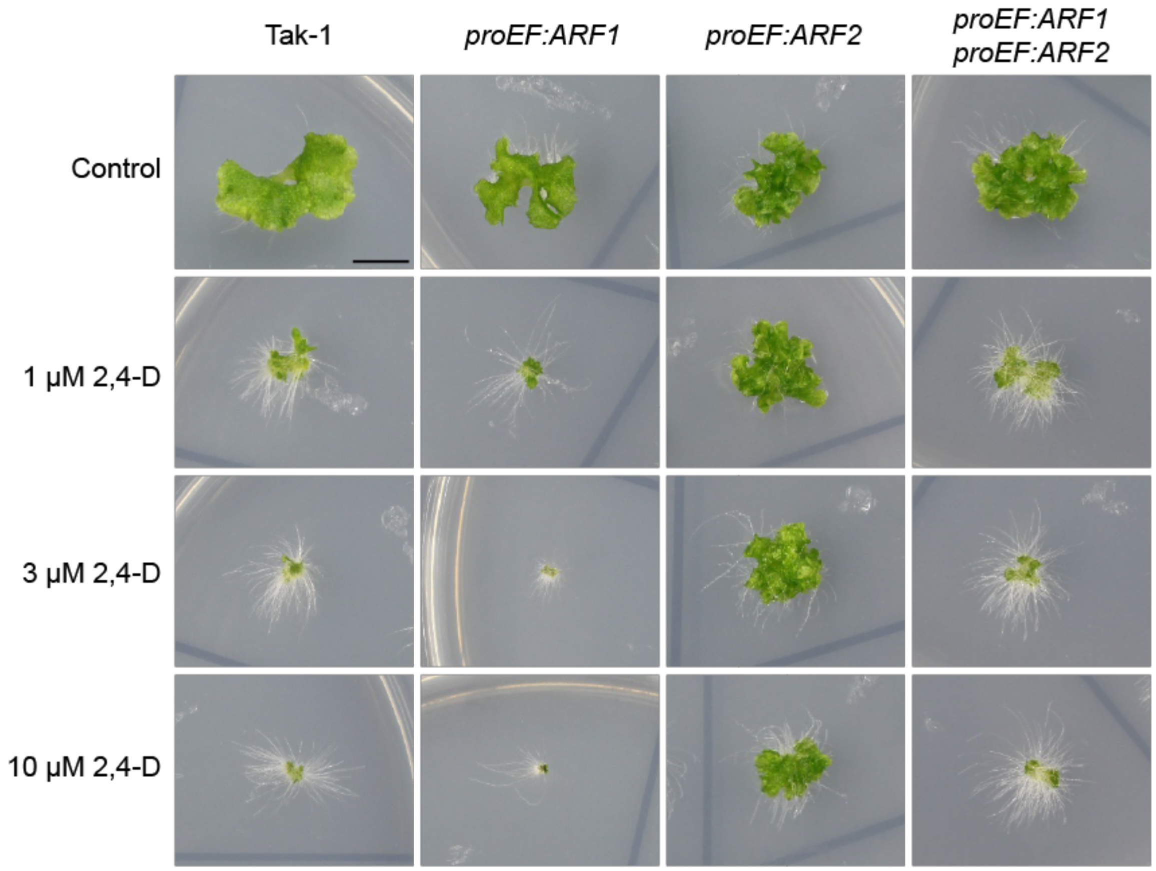


**Fig. S7. Antagonistic action of MpARF1 and MpARF2.**

Fourteen-day-old gemmalings of Tak-1, $_{pro}$*EF:*Mp*ARF1*, $_{pro}$*EF:*Mp*ARF2*, and $_{pro}$*EF:*Mp*ARF1* $_{pro}$*EF:*Mp*ARF2* transgenic lines grown on control medium, or on media containing 1, 3 or 10 µM 2,4-D. The $_{pro}$*EF:*Mp*ARF1* $_{pro}$*EF:*Mp*ARF2* line was generated by introducing the $_{pro}$*EF:*Mp*ARF1* cassette into the presented $_{pro}$*EF:*Mp*ARF2* line. Bar = 1 cm.

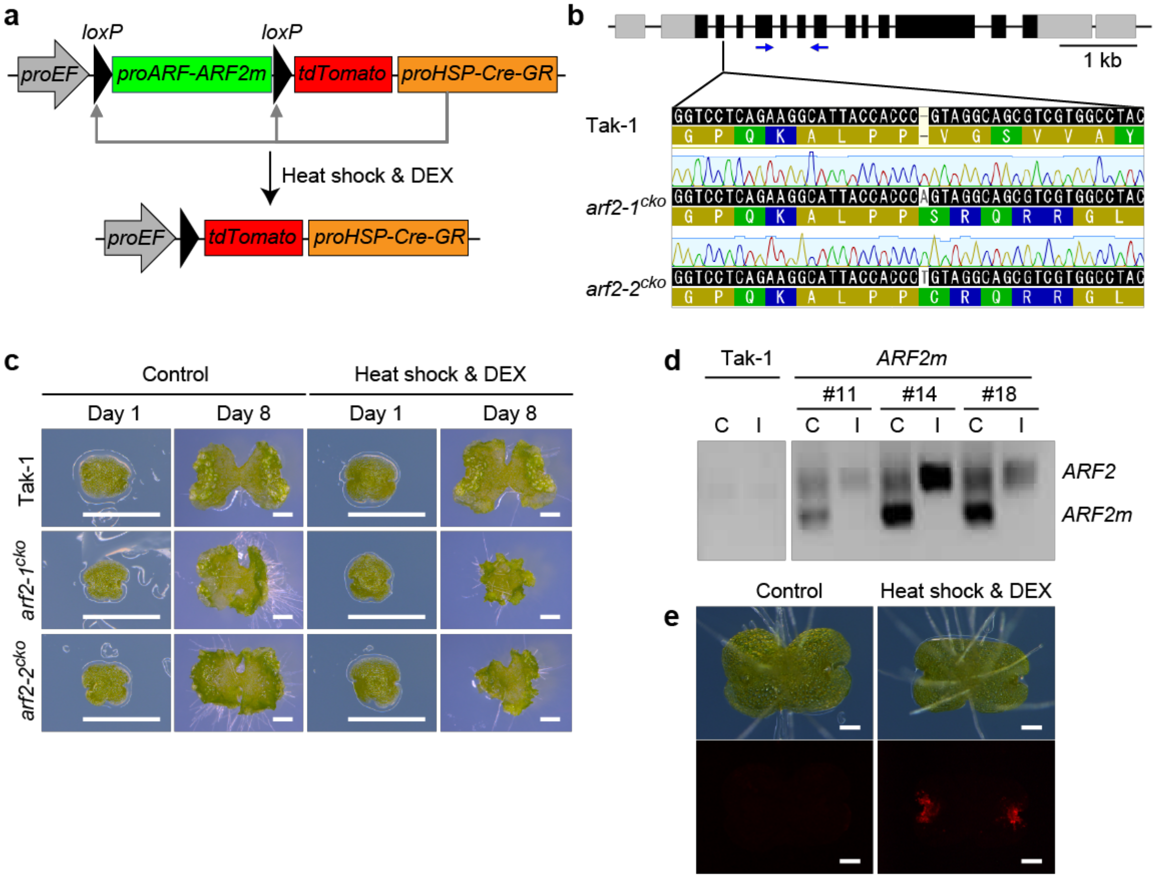


**Fig. S8 Conditional MpARF2 inactivation.**

(**a**) Design of transgene for inducible removal of Mp*ARF2*. A CRISPR-resistant Mp*ARF2* version (*ARF2m*; through engineered silent mutations) is flanked by *loxP* sites and an upstream *EF* promoter and a downstream *tdTomato-NLS* gene, in a construct that also harbors a DEX-inducible Cre-GR version driven from a heat-inducible Mp*HSP17.8A1* promoter ($_{pro}$*HSP*). Following heat shock, and in the presence of DEX, Cre-GR will recombine the two *loxP* sites and excise the *ARF2m* locus. Cells in which recombination occurred are marked by $_{pro}$*EF:tdTomato-NLS* expression. (**b**) CRISPR/Cas9-induced mutations in the endogenous Mp*ARF2* gene in Mp*arf2*$^{cko}$ genetic background. Two different single-base insertions were recovered, generating two independent Mp*arf2*$^{cko}$ alleles. (**c**) Phenotypes of 8-day-old gemmalings of Tak-1, Mp*arf2-1*$^{cko}$ or Mp*arf2-2*$^{cko}$ lines grown on control medium or with 1 µM DEX after a heat shock on day 1. (**d**) PCR amplicons detecting the endogenous *ARF2* (upper band) and *ARF2m* (lower band) genes in Tak-1 and in 3 independent transgenic lines harboring ARF2m (#11, #14, #18; prior to introduction of the CRISPR/Cas9 construct), either grown on control media, or on 1 µM DEX after a heat shock. (**e**) tdTomato expression (lower panels) and appearance (top panels) of 3-day-old Mp*arf2*$^{cko}$ gemmalings grown on control medium or with 1 µM DEX after a heat shock on day 1. Bars are 1 mm in (**c**) and 0.2 mm in (**e**).

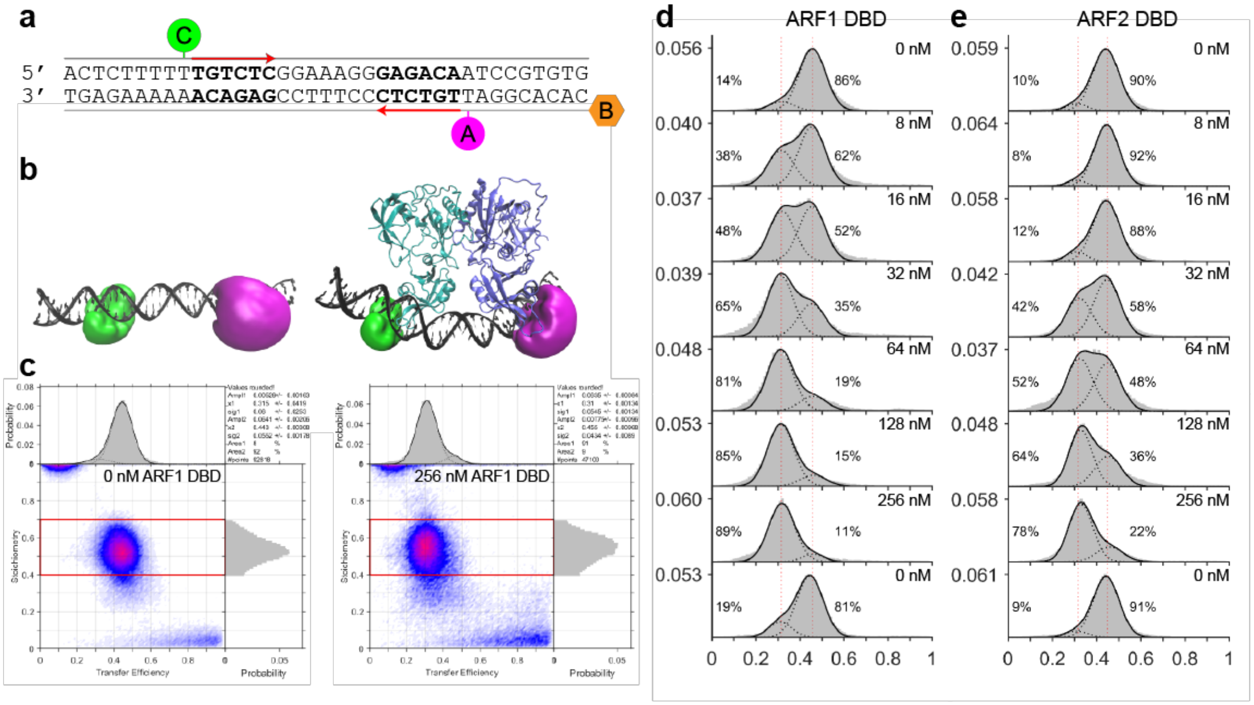


**Fig. S9. Single-molecule FRET analysis of ARF-DNA interaction.**

(**a**) Schematic representation of the ER7 DNA oligonucleotide used in single-molecule FRET experiments, harboring two inverted ARF binding sites (indicated in bold lettering and with a red arrow on top- and bottom-strand, respectively). The bottom-strand was 5’ biotinylated (indicated with an orange hexagon, labeled B) to facilitate immobilization on cover slips. FRET-compatible Cy3B (green circle) and ATTO647N (magenta circle) labels were attached immediately upstream of each ARF binding site indicated with yellow). (**b**) Simulations of accessible volumes of the Cy3B and ATTO647N dyes on the *ER7* oligonucleotide in the absence (left) or presence (right) of MpARF2 DBD (based on the AtARF2-ER7 complex, PDB ID: 6SDG). Note that dye clouds are constrained upon protein binding, predicting reduced FRET efficiency. (**c**) Histograms showing FRET efficiency (x-axis) and Cy3B/ATTO6437N stoichiometry (y-axis) of the immobilized *ER7* oligonucleotide incubated without MpARF1 DBD (left) or with 256 nM MpARF1 DBD (right). Each dot represents a single DNA molecule. Note the shift in FRET efficiency induced by protein binding. (**d**,**e**) Distribution histograms of FRET efficiency (x-axis) of the labeled and immobilized *ER7* oligonucleotide in the presence of increasing concentrations (0-256 nM) of MpARF1 DBD (**d**) or MpARF2 DBD (**e**). Each titration series is followed by an incubation without protein to validate recovery to unbound state. Histograms show fits of the two FRET states representing unbound and bound states, and indicate percentages of DNA molecules in each state.

**Table S1. Crystallography details and refinement statistics (see File S1 for Structure Validation Report)**

| Property | Value |
|---|---|
| PDBid | 6SDG |
| Space group | C 1 2 1 |
| Cell constants (a, b, c (Å), β (º)) | 162.77, 79.36, 79.77, 116.93 |
| Resolution[a] (Å) | 72.6 -2.96 (3.0 – 2.96) |
| Spherical completeness (%) | 73.5 (5.4) |
| Ellipsoidal completeness (%) | 91.7 (30.6) |
| # of reflections | |
| Total | 46775 (170) |
| Unique | 14035 (51) |
| Average multiplicity | 3.3 (3.3) |
| $R_{merge}^{c}$ (%) | 0.09 |
| $R_{sym}^{b}$ (%) | 0.09 |
| <I/σ(I)> | 8.2 (0.7) |
| $CC_{½}$ | 0.996 (0.40) |
| Refinement program | PHENIX 1.16_3549 |
| $R^{d}_{cryst}$, $R^{e}_{free}$ | 0.215, 0.278 |
| $R_{free}$ test set | 768 reflections (5.48%) |
| Wilson B-factor (Å$^2$) | 101.1 |
| Anisotropy | 0.012 |
| Bulk solvent $k_{sol}$(e/Å$^3$), $B_{sol}$ (Å$^2$) | 0.28 , 53.8 |
| Refinement resolution range (Å$^2$), no. reflections | 29.2 - 2.96 (14014) |
| r.m.s. deviation from target values | |
| Bond lengths (Å) | 0.0132 |
| Bond angle distances (Å) | 1.451 |
| Molprobity scores: | |
| Clashscore (‰) | 16.7 |
| Poor rotamers (%) | 3.1 |
| Ramachandran outliers (%) | 0.81 |
| Ramachandran favoured (%) | 93.4 |
| Overall score (‰) | 2.53 |
| Total number of atoms | 5890 |
| Average B, all atoms (Å$^2$) | 112.7 |

[a] Throughout the table, the values in parentheses are for the outermost resolution shell.
[b] $R_{sym} = \Sigma_h \mid \hat{I}_h - I_{h,i} \mid / \Sigma_h\Sigma_i I_{h,i}$ , where $\hat{I}_h = (1/n_h) \Sigma_i I_{h,i}$ and $n_h$ is the number of times a reflection is measured.
[c] $R_{meas} = [\Sigma_h (n_h/[n_h\text{-}1])^{½} \Sigma_i \mid \hat{I}_h - I_{h,i} \mid] / \Sigma_h\Sigma_i I_{h,i}$ , where $\hat{I}_h = (1/n_h) \Sigma_i I_{h,i}$ and $n_h$ is the number of times a reflection is measured.
[d] $R_{cryst} = \Sigma_{hkl} \mid |F_{obs}| - k |F_{calc}| \mid / \Sigma_{hkl} |F_{obs}|$
[e] $R_{free} = \Sigma_{hkl\subset T} \mid |F_{obs}| - k |F_{calc}| \mid / \Sigma_{hkl\subset T} |F_{obs}|$ where T represents a test set comprising ~5% of all reflections excluded during refinement

**Table S2. Oligonucleotides used in this study**

| ID | Sequence |
|---|---|
| ARF1-AAA_F | caccCAGTCAGATCCTCGTAGTCATCTCGCCGCCGCC |
| ARF1-AAA_R | TCACATTTGATCAATGGATACGGCGGCGGCGAGATG |
| ARF1-LFG_F | caccCAGTCAGATCCTCGTAGTCATCTCCTTTTCGGT |
| ARF1-LFG_R | TCACATTTGATCAATGGATACACCGAAAAGGAGATG |
| ARF2-AAA_F | caccGTTCAGAGTGAGCAAAAATGTAAAGCCGCCGCC |
| ARF2-AAA_R | CTATGGCTTGTCGAGAGGCACGGCGGCGGCTTTACA |
| ARF2-IFG_F | caccGTTCAGAGTGAGCAAAAATGTAAAATTTTCGGG |
| ARF2-IFG_R | CTATGGCTTGTCGAGAGGCACCCCGAAAATTTTACA |
| ARF3-AAA_F | caccAAGAGTGCTTCCACTCAGCTGGTGGCCGCCGCC |
| ARF3-AAA_R | TCAGTCGGTGTTGATTGCTTGGGCGGCGGCCACCAG |
| ARF3-IFG_F | caccAAGAGTGCTTCCACTCAGCTGGTGCTGTTCGGC |
| ARF3-IFG_R | TCAGTCGGTGTTGATTGCTTGGCCGAACAGCACCAG |
| HIF | GTATAATGTATGCTATACGAAGTTATGTTT |
| HK009 | CACCATGTATTCTTGTTCGCCG |
| HK011 | GTTTAAACTTAATTAATAACGGCTCAATTTCCCA |
| HK014 | GTTTAAACCGGGGATTCATGGCACCC |
| HK015 | GGCGCGCCTCAGGGGCACCCCCGCTGGGCATC |
| HK018 | CACCATGTCAGAAGCATCTTCC |
| HK020 | GTTTAAACTTAATTAAGAAGGGCTCCACTTCCCA |
| HK021 | TTAATTAACTCTCCCTCCACGACCATC |
| HK022 | GTTTAAACCATTGGATCTCTGGGGACC |
| HK023 | GTTTAAACAAGACGTCGCAAAATTTCC |
| HK024 | GGCGCGCCCTACATGTCGTCGCCGCG |
| HK030 | TTAATTAACTCGACTCTGCCGATGCAA |
| HK031 | GTTTAAACCTCTCTCCTTGTCCAACCC |
| HK032 | GTTTAAACGGTGCCGGAGAGAAGCTCGCCG |
| HK033 | GGCGCGCCCTACTGCGACCGCGTTTT |
| HK064 | GATTGGGAACAGGTTGGAAA |
| HK097 | caccATGCCCGGGCCAAG |
| HK107 | ttttctagaGGTCGGAACTTCTGTCTAAAATGCTAGACT |
| HK108 | caccATGTCAGAAGCATCTTCCATCACTCG |
| HK109 | CATGTCGTCGCCGCGC |
| HK112 | caccATGCCCGGGCCAAGCC |
| HK113 | gtttaaacttaattaaCACCAACTCCACCTGCCAGG |
| HK115 | ttttctagaAAAGAACCGAGGGCAATCGTTC |
| HK116 | ttaattaacACAACCCCTTTCCTGATCTGTCCT |
| HK119 | CCAGCCATCCGCGCTCTC |
| HK120 | gtttaaaccCTGCATAAATTGGCTATCATTTATACTACCATG |
| HK125 | ggttccaatctagatcCCGTCCGAAGATGTGGATGTTG |
| HK184 | ctcgCACGACGCTGCCTACGGG |
| HK185 | aaacCCCGTAGGCAGCGTCGTG |
| HK186 | ctcgTGATACCAGCACGCATGG |
| HK187 | aaacCCATGCGTGCTGGTATCA |

| | |
|---|---|
| HK188 | ctcgCATCCCGTAGTGAGAAGG |
| HK189 | aaacCCTTCTCACTACGGGATG |
| HK190 | CACTGCGGGCAGATTATACC |
| HK201 | AAGCCGTCGAAAAGAAGGAG |
| HK202 | TTCAGGATCGTCCGTTATCC |
| HK279 | tctgtggtcGCCTACTTGCCTCAAGGTCACATAGAG |
| HK280 | acccactccTGGTAATGCCTTCTGAGGACCAGC |
| HK281 | caccCGGGGCCAGAGGAGGACTG |
| HK282 | caccAGAGCTCGCACGGGGAAGTAGTTTAC |
| HK283 | GCGTTTCCTCGTTCTCAAAC |
| HK284 | GGCAGCTTCGACAGACAAG |
| HK285 | tcaCTGCATAAATTGGCTATCATTTATAC |
| HK286 | GGCAGCCAGCCATGTAAGTAG |
| HK287 | CCGGCAGAATTGAGACATTG |
| HK308 | GGTCGAGTGACCTTTGATCG |
| HK309 | GTGGCTGGCTGGATAGTTGG |
| HK481 | tttgtttaaacCCGCAGACGGCTGCTTTTG |
| HK482 | tttggcgcgcctcaCGGTTCCTCAGATAACCCTTCTTGG |
| HK483 | CCTCTCGACAAGCCAACTCCG |
| HK484 | ACATTTTTGCTCACTCTGAACGGGAG |
| HK485 | GCAATCAACACCGACTCCAACAAG |
| HK486 | CAGCTGAGTGGAAGCACTCTTGTTG |
| HK490 | ctagttggaataggttgATGTATTCTTGTTCGCCGATGAGG |
| HK492 | ctagttggaataggttgATGTCAGAAGCATCTTCCATCACTCG |
| HK494 | ctagttggaataggttgATGCCCGGGCCAAGC |
| HK575 | ATCCTCGTCAAATGCCAAACGGAGCTGG |
| HK576 | GGCATTTGACGAGGATCTGACTGTACTTGATCTTGC |
| HK581 | ctagttggaataggttgATGTCATCGTTAAGCAGGGAGCTCG |
| HK582 | agtatggagttgggttgTCTCGGAGCTTGATCAGAATTTTGAATG |
| HK583 | agtatggagttgggttgGGGGCACCCCCGCTG |
| HK584 | agtatggagttgggttgCATGTCGTCGCCGCGC |
| HK585 | agtatggagttgggttgCTGCGACCGCGTTTTGC |
| MpARF2-gt_F1 | CTAAGGTAGCGATTAATGAAAGTGTTCATGCCCGAAG |
| MpARF2-gt_F2 | AACACTAGTGGCGCGGCGTTTCCTCGTTCTCAAAC |
| MpARF2-gt_R1 | CCGGGCAAGCTTTTAATCCACATCTTGGAGCTCCTTT |
| MpARF2-gt_R2 | TTATCCCTAGGCGCGTTCAAAACTCACGCATCGTC |
| MpARF2-L2 | AGGAGGTGGGGCTAAAATGG |
| MpARF2-L6 | GCTCGTCTGTCGTTGCTTTGCTCGTCTGTCGTTGCTTT |
| MpARF2-R1 | ACGTCGTCTTCGTGGTCTGTACGTCGTCTTCGTGGTCTGT |
| MpARF2-R12 | GCTGCCTACGGGTGGTAATGGCTGCCTACGGGTGGTAAT G |
| MpEFp_GT_R1 | GAAGGCTTCTGATTGAAGTTTCCTTTTCTG |
| 21-7_A | TTGTCGGCGATTCGCCGACAA |
| 21-7_B | TTGTCGGCGAATCGCCGACAA |
| GAL4BD_RwithXbaI | TGTTGATAACTCTAGCGATACAGTCAACTG |

| | |
|---|---|
| GAL4BD_FwithXbaI | CACGGGGGACTCTAGAGATCTGAATTCAGGA |
| MpARF1_D-LFG-D_F | caccCAGTCAGATCCTCGTAGTCATCTCCTTTTCGGT |
| MpARF1_D-LFG-D_R | TCACATTTGATCAATGGATACACCGAAAAGGAGATG |
| MpARF1_D-AAA-D_F | caccCAGTCAGATCCTCGTAGTCATCTCGCCGCCGCC |
| MpARF1_D-AAA-D_R | TCACATTTGATCAATGGATACGGCGGCGGCGAGATG |
| MpARF2_S-IFG-D_F | caccGTTCAGAGTGAGCAAAAATGTAAAATTTTCGGG |
| MpARF2_S-IFG-D_R | CTATGGCTTGTCGAGAGGCACCCCGAAAATTTTACA |
| MpARF2_S-AAA-D_F | caccGTTCAGAGTGAGCAAAAATGTAAAGCCGCCGCC |
| MpARF2_S-AAA-D_R | CTATGGCTTGTCGAGAGGCACGGCGGCGGCTTTACA |
| MpARF3_A-LFG-N_F | caccAAGAGTGCTTCCACTCAGCTGGTGCTGTTCGGC |
| MpARF3_A-LFG-N_R | TCAGTCGGTGTTGATTGCTTGGCCGAACAGCACCAG |
| MpARF3_A-AAA-N_F | caccAAGAGTGCTTCCACTCAGCTGGTGGCCGCCGCC |
| MpARF3_A-AAA-N_R | TCAGTCGGTGTTGATTGCTTGGGCGGCGGCCACCAG |
| MpTPL_F | CACCATGTCATCGTTAAGCAGGGA |
| MpTPL_R1 | TCATCTCGGAGCTTGATCAG |
| RenillaLUCforTOPOF | CACCATGACTTCGAAAGTTTATGATCC |
| RenillaLUCstopR | TTATTGTTCATTTTTGAGAACT |
| 6xUASfLUCcloningFforTOPO | CACCTCGAAGCTTGCATGCCTGCAGG |
| 6xUASfLUC_stopR | TTACACGGCGATCTTTCCGCCC |
| ARF1CDS_inf_F1 | GTCGACTGGATCCGGATGTATTCTTGTTCGCCGATGA |
| ARF1CDS_inf_R1 | TAGATATCTCGAGTGGAACGATCGGGGAAATTCG |
| MpARF2_c1f_TOPO | CACCATGTCAGAAGCATCTTCCATC |
| MpARF2_stop | CATGTCGTCGCCGCGC |
| SJD005 | gcgattaattaagctTTGGCAGAATGAGTGTGGCCAATTCTC |
| SJD006 | TGCCGCCGCCaagcttGGGGCACCCCCGCTGGG |
| SJD007 | taaactagtggcgcgATCTCGTTTTCACACTCCACACTCCGAT |
| SJD008 | ttatccctaggcgcgTTCCTCCCCCGCTCGAAATCAC |
| SJD009 | gcgattaattaagctTGAGCGAGCAACACGAAGACTC |
| SJD010 | TGCCGCCGCCaagcttCATGTCGTCGCCGCGCGC |
| SJD011 | taaactagtggcgcgACCATATGCCTTTTTTATGTATTTAAATTCTTCATCTATTCTC |
| SJD012 | ttatccctaggcgcgACTCCGCAATTTCGCCTCAGAGTG |
| HK056 | TGGCCCAGTTCACCAACTAT |
| SJD091 | CATCGGAGTGTGGAGTGTGA |
| HK072 | GATCGATCGACGTCCAAAGT |
| SJD094 | GGCGAGACGGAAGAACAGAG |
| SJD149 | TGGAGCCGTACATGAACTGAGG |
| HK050 | CATCTTCTCACAACTGGATGGA |
| HK063 | GCTCCATCCTTCGCAAGC |
| MpTPL-mNeongreen FW | gctgactctagcagatcATGTCATCGTTAAGCAGGGAG |
| MpTPL-mNeongreen RV | GCCCGGTACCagaTCTCGGAGCTTGATCAGAAT |
| MpARF1-mScarlet-I FW | gctgactctagcagatcATGTATTCTTGTTCGCCCGATG |
| MpARF1-mScarlet-I FW | GCCCGGTACCagaGGGGCACCCCCGCTGG |
| MpARF2-mScarlet-I FW | gctgactctagcagatcATGTCAGAAGCATCTTCCATC |

| | |
|---|---|
| MpARF2-mScarlet-I FW | GCCCGGTACCagaCATGTCGTCGCCGCGCGCCCC |
| MpARF1-DBD FW | tttaagaaggagaattcATGTATTCTTGTTCGCCGATGAGG |
| MpARF1-DBD RV | attgattgggagtaagaattcccCGGCTCAATTTCCCACAGAGAC |
| MpARF2-DBD FW | tttaagaaggagaattcATGGCGAGCAGACAGCTTGC |
| MpARF2-DBD RV | attgattgggagtaagaattcccGGGCTCCACTTCCCATGGTGAGACC |
